# Selective Control of Synaptically-Connected Circuit Elements by All-Optical Synapses

**DOI:** 10.1101/2021.10.29.466531

**Authors:** Mansi Prakash, Jeremy Murphy, Robyn St Laurent, Nina Friedman, Emmanuel L. Crespo, Andreas Bjorefeldt, Akash Pal, Yuvraj Bhagat, Julie A. Kauer, Nathan C. Shaner, Diane Lipscombe, Christopher I. Moore, Ute Hochgeschwender

## Abstract

Understanding percepts, engrams and actions requires methods for selectively modulating synaptic communication between specific subsets of interconnected cells. Here, we develop an approach to control synaptically connected elements using bioluminescent light: Luciferase-generated light, originating from a presynaptic axon terminal, modulates an opsin in its postsynaptic target. Vesicular-localized luciferase is released into the synaptic cleft in response to presynaptic activity, creating a real-time ‘Optical Synapse’. Light production is under experimenter-control by introduction of the small molecule luciferin. Signal transmission across this optical synapse is temporally defined by the presence of both the luciferin and presynaptic activity. We validate synaptic ‘Interluminescence’ by multi-electrode recording in cultured neurons and in mice *in vivo*. Interluminescence represents a powerful approach to achieve synapse-specific and activity-dependent circuit control during behavior *in vivo*.

## Introduction

A wealth of new tools are revolutionizing neuroscience by allowing direct control of specific subpopulations of neurons for brief and sustained time periods (e.g., opto-, chemo- and sonogenetics;^1–3^). The ability to regulate genetically identified neurons in selected brain areas has been a significant benefit to studying neural dynamics and its link to behavior.

However, information processing leading to percepts, memories and/or actions requires multiple nodes acting in sequence within a network, with multiple cell types in multiple areas conducting specific cell-to-cell communication. The ideal tool(s) in the next generation of approaches will allow intersectional circuit dissection—specifying and regulating participants at multiple stations. Further, tools for fully understanding systems underlying behavior will allow them to demonstrate natural activity, enhancing or suppressing, for example, transmission of endogenous patterns. While real-time feedback interventions driven by computer recognition of activity patterns are increasingly being applied to electrical stimulation (e.g., deep-brain stimulation) systems, most tools that have genetic precision and molecular specificity are still regulated *en masse* by rising gradients of sustained chemical drivers or by imposed patterns of optogenetic drive. Tools that permit direct experimental control of the efficacy and form of synaptic transmission between specific partners will be a key step in providing the next wave of insight into network dynamics and function.

The most common strategy currently in use to modulate specific synaptic connections (**Fig. 1a**) involves light-activation of opsin-expressing presynaptic neurons using localized fiber optic stimulation near the postsynaptic target neurons (**Fig. 1b**^4^). A high degree of presynaptic specificity can be achieved by this approach, using retrograde and Cre-dependent expression of optogenetic elements and localizing the light source, yet postsynaptic specificity will depend on the extent of presynaptic contact spread. A conceptually similar approach is achieved using chemogenetic methods to modulate synaptic transmission by targeting axon terminals (**Fig. 1c**). Chemogenetic neuromodulation can be restricted to a subset of target neurons, leaving other interconnected areas unaffected. For example, DREADDs expressed in long-range projecting neurons in one cortical layer can be activated by CNO ligand application to a confined area in another cortical layer^5^. These methods are limited by studying cell-cell communication between populations that are anatomically separate because 1 mm^3^ of light (optogenetic) or exogenous drug (chemogenetic) will likely act on multiple cells. Further, most neural computations take place between highly interspersed cells in tightly packed spaces.

**Fig. 1.**
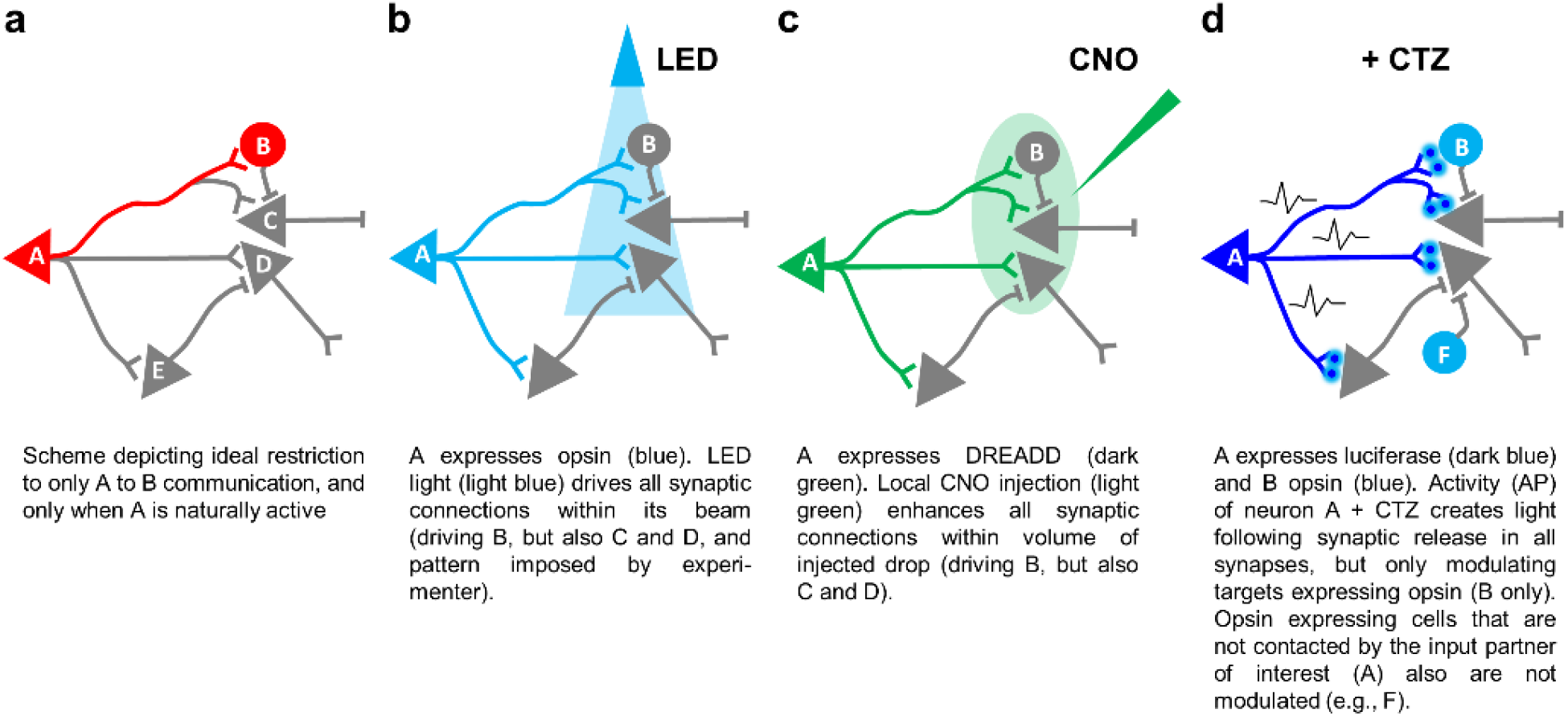
Transsynaptic modulation. **a** Cell A connects with cells B, C, D and E; however, only cell A’s communication with cell B should be modulated, either amplified or dampened. **b** If cell A expresses optogenetic actuators, restriction of a light beam to the area of intended synaptic transmission can minimize unwanted activation (cell E will not be activated), but the likelihood of still activating unwanted synapses (cells C and D) is high. **c** The same applies when expressing a chemogenetic actuator in cell A and restricting application of the ligand CNO *(Clozapine-N-* Oxide) to an anatomical area as small as possible. **d** True cell-to-cell synaptic communication can be achieved by expressing a luciferase in cell A and an opsin in cell B. Activity of neuron A and application of the luciferin CTZ (Coelenterazine) results in light emission at all synapses of A, but only the opsin expressing cell B will be modulated. At the same time, opsin expressing cells not synaptically contacted by the luciferase expressing cell A will not be modulated.

Here, we describe the Optical Synapse, an approach to control synaptic connectivity that utilizes presynaptically originating BioLuminescence (BL) to activate OptoGenetic (OG) actuators expressed at postsynaptic sites (**Fig. 1d**). We have shown that BL-OG can operate within a cell, with a luciferase tethered to an opsin (a ‘Luminopsin’)^6–9^. Photon release from the luciferin-luciferase interaction activates the associated opsin thereby achieving optogenetic modulation. Depending on the biophysical properties of the opsin, excitatory or inhibitory, bioluminescence may depolarize or hyperpolarize the neuron.

In our *Interluminescent* Optical Synapse, we used BL-OG to achieve synapse-specific and activity-dependent circuit control, by expressing the luciferase presynaptically and its partner opsin postsynaptically. When presynaptic luciferase and postsynaptic opsin are present at the same synapse, BL-OG modulation is triggered when luciferin is provided, a requirement that allows experimental control of intersectional communication. The spatial requirements of Interluminescence in the current application restricts it to synapses that express the luciferase in presynaptic vesicles and opsins postsynaptically. The release of luciferase from the presynaptic terminal depends on presynaptic depolarization, similar to synaptic transmission via neurotransmitters or neuropeptides. Here we describe examples of the transmission of bioluminescence signals across synapses in culture and *in vivo*, and through a series of independent tests show that the postsynaptic output is mediated by optical coupling, and independent of classic neurotransmitter-mediated synaptic transmission. This new intersectional technology can provide a novel class of cell-pairing and activity-specific control for testing the mechanisms underlying behavior.

## Results

### Interluminescence via an Optical Synapse

We used cultured cortical neurons to first establish if a luciferase genetically targeted to presynaptic vesicles could be released into the synaptic cleft and generate sufficient photon density to activate opsins genetically targeted to a postsynaptic cell. We term this form of bioluminescence-mediated synaptic transmission Interluminescence (**Fig. 2a**). Critically, we needed to show that Interluminescence: i) generates a measurable postsynaptic response; ii) mediates different postsynaptic responses depending on the type of opsin, with cation permeable opsins triggering postsynaptic depolarization and excitation, and anion permeable opsins hyperpolarization and inhibition; iii) occurs when luciferase is co-released with endogenous transmitters and peptides, but only in the presence of the luciferin; iv) occurs independent of classic neurotransmission; and, v) can co-exist with classic neurotransmission.

**Fig. 2.**
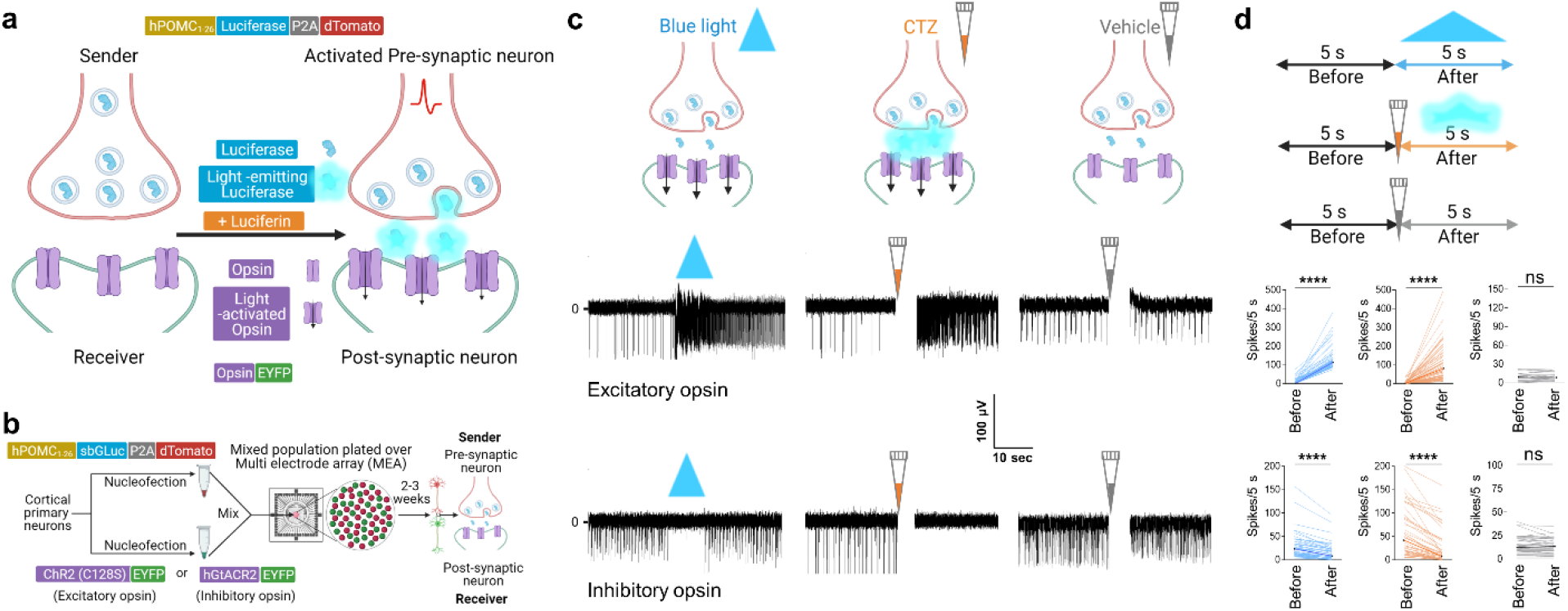
Modulation of postsynaptic neural activity by Interluminescence in mixed cultures of pre- and postsynaptic neurons. **a** Schematic: Interluminescence via an Optical Synapse. Luciferases (blue colored enzyme inside the grey circles) are released from presynaptic vesicles and, in the presence of the luciferin, emit light (bioluminescence: light bluish glow) that activates postsynaptic opsins (magenta; downward black arrows indicating ion movement through open channels). **b** Schematic of experimental design and constructs used for separate nucleofections of cortical neurons with either hPOMC1-26-sbGLuc-P2A-dTomato or one of the opsins, ChR2(C128S)-EYFP (excitatory) and hGtACR2-EYFP (inhibitory); neurons were then mixed (red and green spheres), plated on MEAs and maintained for 2-3 weeks until recording. **c** Illustrations (upper panels) and corresponding representative traces from individual electrodes of MEAs (middle and lower panels) showing response of postsynaptic neurons to blue light (blue solid triangle), CTZ-induced bioluminescence (orange pipette tip) and vehicle (gray pipette tip) expressing the excitatory opsin ChR2(C128S) (middle panels) and the inhibitory opsin hGtACR2 (lower panels). **d** Schematic showing the time windows for analyzing the number of spikes for each treatment (upper panel) and ladder plots from multiple experiments (middle panels for excitatory and lower panels for inhibitory opsin expressing postsynaptic neurons as depicted for individual traces in (c). ChR2(C128S), blue light, n=62, p<0.0001, CTZ, n=62, p<0.0001, vehicle, n=24, p=0.6498; hGtACR2, blue light, n=49, p<0.0001, CTZ, n=49, p<0.0001, vehicle, n=49, p=0.5594; Wilcoxon matched-pairs signed rank test. The artifacts due to addition of reagents in MEAs are overlaid by a vertical white bar in all MEA recording traces (the white gap right after addition of either CTZ or vehicle). ns, not significant; ****, p<0.0001

### Interluminescence modulates spontaneous neural activity in culture

We targeted the blue light emitting luciferase sbGLuc, a bright *Gaussia* luciferase variant^10^, to synaptic vesicles in cortical neurons using the vesicle targeting sequence of the human pro-opiomelanocortin pro-peptide (hPOMC1-26)^11,12^. The targeting construct also contained the reporter gene dTomato attached to sbGLuc via a P2A cleavage sequence (**Fig. 2a**). In addition to being a bright photon source, sbGLuc is favorable because it is also stable at the lower pH levels of synaptic vesicles^13,14^.

We selected high light sensitivity opsins for our initial experiments. We employed the excitatory step function opsin ChR2(C128S)^15^ and the inhibitory anion channel hGtACR2^16^ as both exhibit high sensitivity to blue light relative to other opsins. In our initial studies, we separately nucleofected cortical neurons with hPOMC1-26-sbGLuc, ChR2(C128S), or hGtACR2 to ensure that luciferase and opsin were expressed in different cell populations (**Fig. 2b**). We then mixed the two populations of cells, one luciferase expressing and one opsin expressing, and plated the mixture on multi electrode arrays (MEAs)^17^. We used externally presented blue light to activate opsins directly and showed that this increased (**Fig. 2c** middle panel; ChR2(C128S)) or decreased (**Fig. 2c**, lower panel; hGtACR2) the activity of the culture, as expected based on opsin expressed. We then added the luciferin Coelenterazine (CTZ), the substrate for *Gaussia* luciferases, and observed increased (ChR2(C128S)) or decreased (hGtACR2) spontaneous activity consistent with the expressed opsin (**Fig. 2c**). By contrast, the addition of the vehicle alone had no consistent impact on ongoing neural activity.

We used direct LED stimulation of the postsynaptic opsins as an internal control and compared the responses of cultures to LED (blue light), bioluminescence (CTZ), and control (vehicle) in three independent experiments (summarized in the ladder plots in **Fig. 2d**). Spontaneous activity levels varied across microelectrode arrays (MEAs) as is typical (see also **Supplementary Fig. 1**), so we assessed the effect of each manipulation by comparing spike rate over time before and after treatment from each electrode and across different MEAs. We observed a significant difference in spontaneous activity in cultures before and after stimulation by blue light LED and CTZ but no consistent differences in neural activity before and after vehicle (ChR2(C128S), blue light, n=62, p<0.0001, CTZ, n=62, p<0.0001, vehicle, n=24, p=0.6498; hGtACR2, blue light, n=49, p<0.0001, CTZ, n=49, p<0.0001, vehicle, n=49, p=0.5594; Wilcoxon matched-pairs signed rank test). Our data also show that the activity of opsin expressing postsynaptic neurons could be augmented or inhibited following presynaptic activation in the presence of CTZ depending on the nature of the postsynaptic opsin.

### Interluminescence requires connections between pre- and postsynaptic neurons

To assess the properties of CTZ-dependent responses in more detail, and directly test if synaptic connectivity is required, we employed a co-culture system in which presynaptic and postsynaptic populations are seeded in separate compartments and then allowed to form synapses across a separating gap (**Fig. 3a**). We nucleofected ‘presynaptic’ cortical neurons with luciferase, and ‘postsynaptic’ hippocampal or striatal target neurons with ChR2(C128S).

**Fig. 3.**
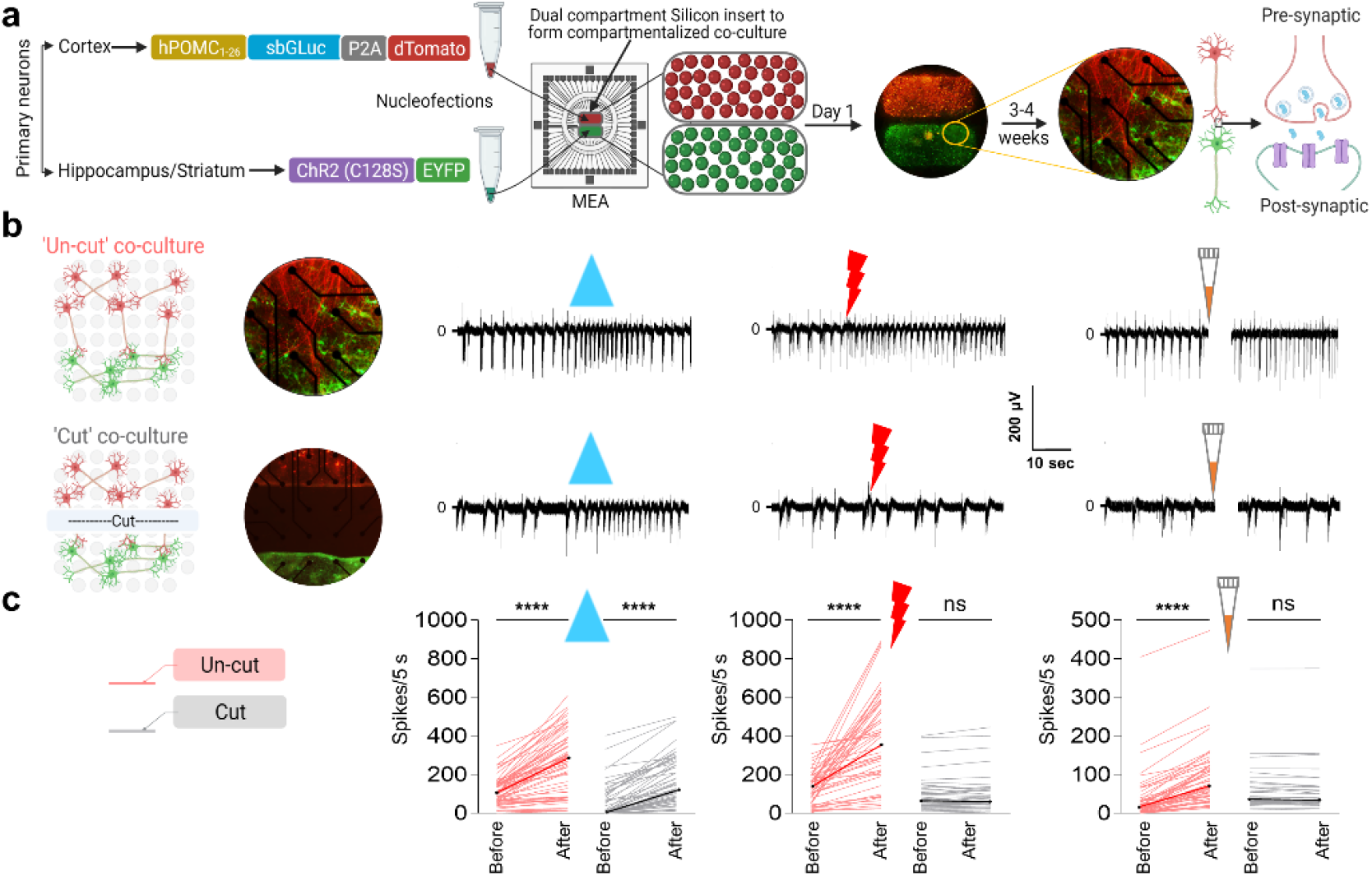
Communication via Interluminescence in co-cultured populations depends on intact synaptic connections. **a** Schematic of experimental design and constructs used for separate nucleofections of cortical and hippocampal or striatal primary neurons. Cortical neurons were nucleofected with the luciferase construct (hPOMC1-26-sbGLuc-P2A-dTomato) and plated in the upper compartment of a 2-chamber silicon divider (group of red spheres covering the upper half of MEA electrodes), and their natural synaptic targets, hippocampal or striatal neurons, were nucleofected with the excitatory opsin construct (ChR2(C128S)-EYFP) and plated in the lower compartment (group of green spheres covering the lower half of MEA electrodes). The next day neurons had attached and the divider was removed (fluorescent image; 5 x magnification) showing the expression of luciferase and opsin in respective neuronal populations). Cultures matured over the next 3 weeks, with processes from cortical neurons growing deep into the hippocampal or striatal areas (fluorescent image; 20x magnification) showing the processes from the cortical population (dTomato) contacting the hippocampal neurons (EYFP). **b** Illustrations showing the layout of electrodes (light grey circles) in 1-well MEAs with a co-culture (left panel) when the inter-population connections are intact (‘un-cut’: upper left) or severed (‘cut’: lower left) by making a ‘cut’ between the two populations. Recordings from one electrode within the postsynaptic population (right panels) when treated with blue light (blue solid triangle), pre-synaptic electrical stimulation (red bolt), and CTZ (orange pipette tip) with connecting processes were intact (right upper) versus cut (right lower). **c** Schematic showing the color code (left panel) used for the ladder plots for both ‘un-cut’ (peach) and ‘cut’ (gray) co-cultures in the right panel. Ladder plots (right panel) showing change in number of spikes of post-synaptic neurons 5 seconds before and after each treatment (blue light; electrical stimulation; CTZ) for both the un-cut (peach) and ‘cut’ (gray) conditions (blue light, un-cut n=63, p<0.0001, cut n=58, p<0.0001; electrical stimulation, un-cut n=54, p<0.0001, cut n=32, p=0.0965; CTZ, un-cut n=51, p<0.0001, cut n=38, p=0.7388; Wilcoxon matched-pairs signed rank test). The artifacts due to addition of reagents in MEAs are overlaid by a vertical white bar in all MEA recording traces (the white gap right after addition of CTZ). ns, not significant; ****, p<0.0001

We observed an increase in spontaneous activity and in spiking synchrony within the MEAs as the cultures matured (**Supplementary Fig. 2**). The physical separation of pre- and post-synaptic neurons allowed us to know which population was active in different treatment conditions (**Supplementary Fig. 3**). Direct electrical stimulation of presynaptic neurons increased spiking in ChR2(C128S) expressing postsynaptic neurons presumably through classic synaptic neurotransmission (**Supplementary Fig. 3c**). Blue light LED stimulation (which directly activates the postsynaptic Opsin) or CTZ addition (which depends on ongoing spontaneous presynaptic activity) by contrast, increased spiking in ChR2(C128S) expressing postsynaptic neurons but without a change in the activity of cortical presynaptic neurons (**Supplementary Fig. 3d,e**). We also showed that only electrodes on the MEA that responded to direct optogenetic stimulation (LED) responded to CTZ application, an independent demonstration of the specificity of CTZ-dependent responses.

We next tested whether CTZ-dependent changes in postsynaptic neural activity were mediated by synaptic events and presynaptic depolarization. We compared responses before and after severing the connections between the pre- and postsynaptic populations (**Fig. 3b**, upper versus lower panels). As expected, LED stimulation, which activates postsynaptic opsins directly, induced increased postsynaptic spiking equally in ‘un-cut’ and ‘cut’ co-cultures. In contrast, direct electrical stimulation of presynaptic neurons and CTZ-induced bioluminescence failed to alter the excitability of postsynaptic neurons in separated co-cultures. This result is consistent with the hypothesis that CTZ modulation of postsynaptic activity requires synaptic connectivity. The results from several independent experiments of presynaptic (cortical) and postsynaptic (hippocampal, striatal) neural cultures are summarized (**Fig. 3c**; blue light, un-cut n=63, p<0.0001, cut n=58, p<0.0001; electrical stimulation, un-cut n=54, p<0.0001, cut n=32, p=0.0965; CTZ, un-cut n=51, p<0.0001, cut n=38, p=0.7388; Wilcoxon matched-pairs signed rank test).

In sum, these data provide strong support for Interluminescence as a form of engineered synaptic transmission that achieves cell-cell communication via bioluminescence originating presynaptically activating opsins postsynaptically.

### Interluminescence depends on presynaptic activity and occurs independent of classic synaptic neurotransmission

We found that addition of a cocktail of transmitter receptor blockers that included NBQX, a selective blocker of non-NMDA mediated synaptic transmission, D-AP5, a NMDA receptor antagonist, Gabazine and CGP55845, antagonists at GABAA and GABAB receptors, respectively, and Strychnine, a glycine receptor antagonist, silenced spontaneous activity of the entire neural culture on MEAs for several minutes (**Supplementary Fig. 4a, b(i), c**; n=27, P<0.0001; Mann-Whitney test) but did not impact direct activation of postsynaptic opsin-expressing neurons by blue light LED (**Supplementary Fig. 4b(ii), c**; n=27, P<0.0001; Mann-Whitney test). We employed this neurotransmission blockade cocktail to test the requirements for bioluminescence-mediated synaptic transmission separated from endogenous transmitter mediated effects (**Fig. 4**). When CTZ was delivered with, or immediately after, addition of blockers, postsynaptic activation through bioluminescence was robust, while vehicle addition had no effect (**Fig. 4b,c**; c(i): SB alone, n=27, c(ii): SB + CTZ, n=37, c(iii): SB + vehicle, n=38; SB alone (after) v/s SB + CTZ (after) p<0.0001; SB + CTZ (after) v/s SB + vehicle (after) p<0.0001; SB alone (after) v/s SB + vehicle (after) P=0.7305; Mann-Whitney test). However, addition of CTZ more than 20 seconds after blockers had no effect on spiking of opsin expressing neurons (**Fig. 4d(i), e(i)**: n=18, SB (after) v/s CTZ added later (after) p=0.4022; Mann-Whitney test).

**Fig. 4.**
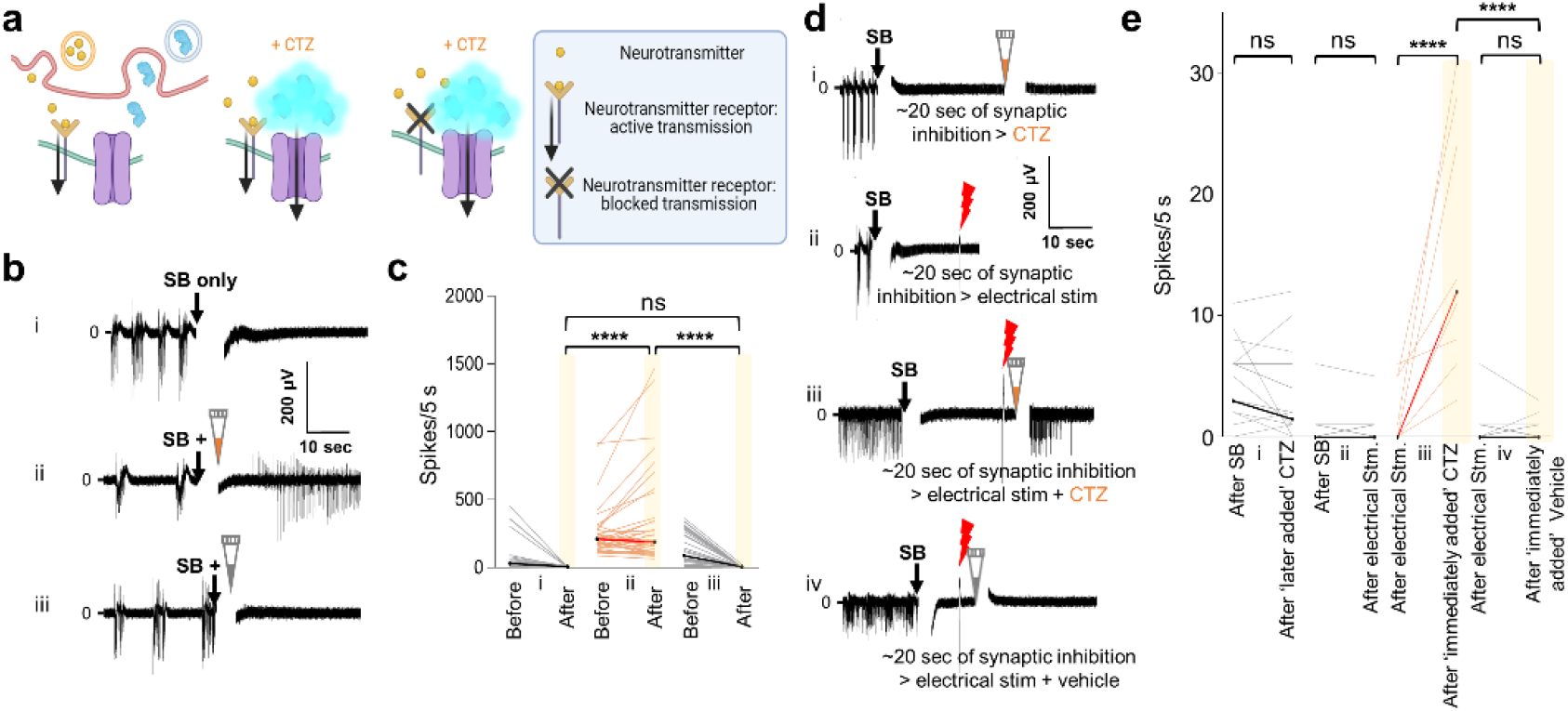
Interluminescence elicits postsynaptic firing increase in the presence of synaptic blockers dependent on presynaptic neuronal activity. **a** Illustrations showing release of synaptic vesicle contents (neurotransmitters: yellow spheres, luciferases: blue enzymes) with spontaneous presynaptic activity inducing postsynaptic responses with transmitters alone (left panel), with transmitters and bioluminescent activation of opsins in the presence of CTZ (middle panel), and the effect of application of synaptic blockers (SB), allowing to isolate the effects of bioluminescence-mediated synaptic transmission (right panel). **b** Traces from representative electrodes of opsin expressing population applying to the culture (i) synaptic blockers alone, (ii) synaptic blockers together with CTZ, or (iii) synaptic blockers together with vehicle. **c** Ladder plots of recordings under the conditions depicted in (b) from electrodes across opsin expressing populations comparing number of spikes 5 seconds before and after (i) synaptic blockers alone, (ii) synaptic blockers together with CTZ, or (iii) synaptic blockers together with vehicle. (i) SB alone, n=27, (ii) SB + CTZ, n=37, (iii) SB + vehicle, n=38; SB alone (after) v/s SB + CTZ (after), p<0.0001; SB + CTZ (after) v/s SB + vehicle (after), p<0.0001; SB alone (after) v/s SB + vehicle (after), P=0.7305; Mann-Whitney test. **d** Traces from representative electrode recordings of opsin expressing population applying to the culture synaptic blockers followed after ~20 seconds by application of (i) CTZ, (ii) electrical stimulation, and electrical stimulation together with either (iii) CTZ or (iv) vehicle. **e** Ladder plots of recordings under the conditions depicted in (d) across populations. (i), n=18, SB (after) v/s CTZ added ~20s later (after), p=0.4022; (ii), n=35, SB (after) v/s electrical stimulation (after) p>0.9999; (iii), n= 10; electrical stimulation (after) v/s immediate CTZ treatment (after) (first yellow bar), p<0.0001; (iv), n=35, electrical stimulation (after) v/s immediate vehicle treatment (after), p=0.8553; Mann-Whitney test. Significant increase in activity of opsin expressing populations is observed only when CTZ (first yellow bar) and not when vehicle (second yellow bar) is applied immediately following electrical stimulation (immediate CTZ (after), (n=10) v/s immediate vehicle (after), (n=35), p<0.0001; Mann-Whitney test. The artifacts due to addition of reagents in MEAs are overlaid by a vertical white bar in the recording traces (the white gap right after addition of SB, CTZ or vehicle). Artifacts due to electrical stimulation are visible under the red bolts. ns, not significant; ****, p<0.0001

This finding supports the conclusion that in a spontaneously active culture sufficient luciferase concentration is present in the synaptic cleft right after its release, such that CTZ addition immediately after acute inhibition of all neural activity will create photon density great enough to enable opsin activation. In contrast, once spontaneous activity of the culture has stopped for at least 20 seconds, addition of CTZ has no effect as luciferases have diffused away from the cleft, dropping photon density below a level that allows opsin activation. This interpretation is further confirmed by CTZ induction of spiking during the ‘silent’ phase with preceding electrical stimulation that acutely releases luciferases into the cleft (**Fig. 4d,e**; e(iii): n=10, electrical stimulation (after) v/s immediately added CTZ (after) p<0.0001; Mann-Whitney test), while blockers (after) with electrical stimulation alone (after) or electrical stimulation (after) with immediate vehicle addition (after) had no effect (**Fig. 4d,e**; e(ii): n=35, p=0.9999; e(iv): n=35, p=0.8553; Mann-Whitney test). Significant increase in activity of opsin-expressing populations is observed only when CTZ, and not vehicle, is applied immediately after electrical stimulation (**Fig. 4e**; first yellow bar v/s second yellow bar; immediate CTZ (after), n=10, v/s immediate vehicle (after), n=35, p<0.0001; Mann-Whitney test). These findings show that Interluminescence depends on presynaptic activity and occurs independent of the postsynaptic action of classic synaptic neurotransmitters and neurotransmitter receptors.

### Interluminescence depends on presynaptic vesicle release

To test whether presynaptic vesicle fusion is essential for Interluminescence, we used botulinum toxin (BoNT) to inhibit this process. BoNT, a neurotoxin that cleaves SNARE proteins, inhibits vesicular fusion and cargo release from presynaptic vesicles^18^ (**Fig. 5a**). BoNT-treated cultures showed decreased luciferase concentration in the media, as expected from decreased vesicle release (**Supplementary Fig. 5**). When presynaptic neurons were electrically stimulated and CTZ added immediately after, Interluminescence-induced spiking was not observed in cultures treated with BoNT, consistent with inhibition of synaptic vesicle fusion and block of luciferase release into the synaptic cleft (**Fig. 5b, c** for individual traces and ladder plots; n=21, electrical stimulation + CTZ p=0.7173; Wilcoxon matched-pairs signed rank test). In contrast, direct application of blue light elicited robust spiking in opsin-expressing postsynaptic neurons (n=21, blue light LED p<0.0001; Wilcoxon matched-pairs signed rank test). BoNT inhibits exocytosis of small and large dense core synaptic vesicles (LDCVs) that contain small molecule neurotransmitters and peptides, respectively. We used the sorting signal for a neuropeptide, POMC, to concentrate luciferase in presynaptic vesicles using an hPOMC1-26-sbGLuc-eGFP fusion protein. We assessed localization of sbGLuc-eGFP utilizing an antibody to dopamine β-hydroxylase which labels LDCVs^19^ (**Supplementary Fig. 6**). We observed partial co-localization of eGFP and anti-dopamine β-hydroxylase, confirming the presence of sbGLuc in dopamine β-hydroxylase expressing synaptic vesicles, but we also observed sbGLuc-eGFP signals consistent with a considerable fraction of luciferases in other vesicles.

**Fig. 5.**
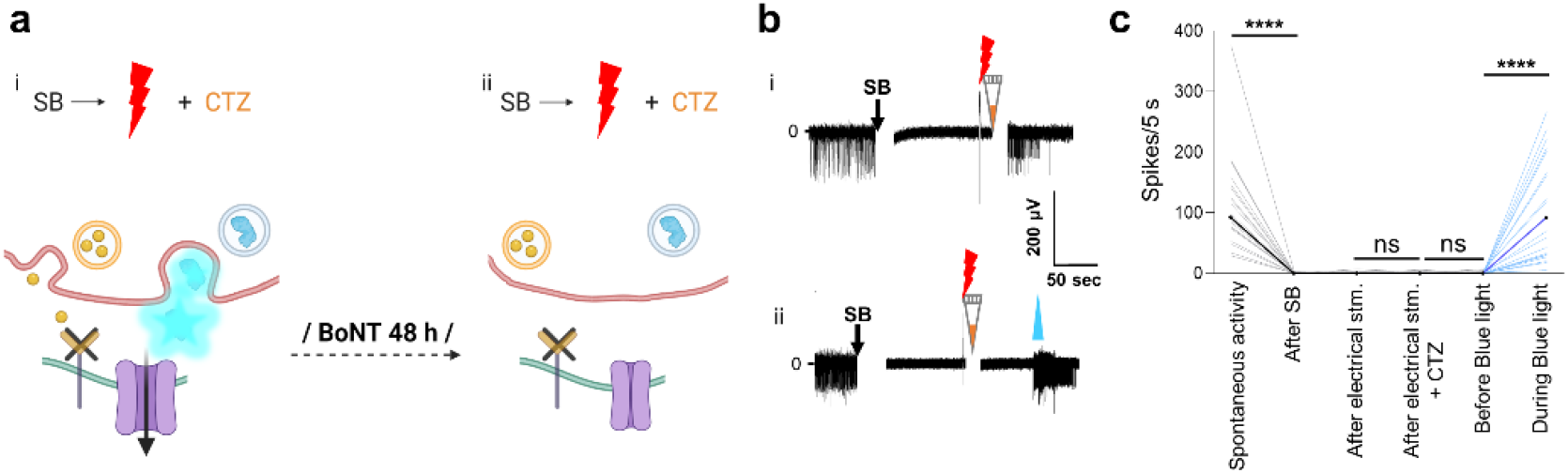
Interluminescence is dependent on presynaptic vesicle release. **a** Schematics of synapses receiving synaptic blockers (SB) followed by electrical stimulation of presynaptic neurons together with CTZ application without (i) and after 48 h Botulinum Neurotoxin (BoNT) treatment (ii). **b** Representative trace of MEA recordings of opsin expressing neurons without BoNT treatment (i, from Fig. 4diii) and 48 h after BoNT treatment of the co-culture (ii): electrical stimulation of pre-synaptic neurons together with CTZ application fails to elicit firing after BoNT treatment, while blue light still induces firing in the same recording. **c** Ladder plots of recording conditions as in (b) from electrodes across populations (comparisons are: spontaneous activity vs SB addition (after), n=21, p<0.0001; electrical stimulation (after) vs immediately following CTZ (after), n=21, p=0.7173; electrical stimulation + CTZ (after) vs blue light (before), n=21, p=0.6055; blue light (before) vs blue light (during), n=21, p<0.0001; Wilcoxon matched-pairs signed rank test). The artifacts due to addition of reagents in MEAs are overlaid by a vertical white bar in the recording traces (the white gap right after addition of SB or CTZ). Artifacts due to electrical stimulation are visible under the red bolts. ns, not significant; ****, p<0.0001

### Interluminescence-mediated activation of postsynaptic neurons requires opsins

We performed two experiments to test if bioluminescence-mediated activation of postsynaptic neurons occurs through opsins. First, we inactivated the step function opsin ChR2(C128S) by exposure to longer wavelength light (590 nm). This protocol resulted in complete opsin inactivation (**Fig. 6a** schematic, **6b** upper trace). To test if this inactivation also blocked Interluminescence, we applied blockers and CTZ to spontaneously spiking neurons, resulting in a neurotransmitter independent increase in spiking through bioluminescent activation of the opsin (**Fig. 6b**, lower trace). 590 nm light completely abolished spiking initiated by CTZ (**Fig. 6b**, lower trace) consistent with a need for recruitable opsins for Interluminescence (**Fig. 6c** ladder plot; SB + CTZ, n=51, p=0.0369; Green Light, n=51, p=0.0001; Wilcoxon matched-pairs signed rank test). Second, we used a non-functional opsin mutant ChR2(C128S)-E97R-D253A that does not produce photocurrent^20^ (**Fig. 6d** schematic, **Fig. 6e** upper trace). In cultures expressing inactive ChR2(C128S) in postsynaptic neurons, CTZ generated bioluminescence, but no increase in spiking (**Fig. 6e**, lower trace, **6f** ladder plot; SB alone (after), n=21, v/s SB + CTZ (after), n= 49, p=0.7870; Mann-Whitney test). These data indicate that Interluminescence is mediated by photocurrent generation following bioluminescent activation of the opsin.

**Fig. 6.**
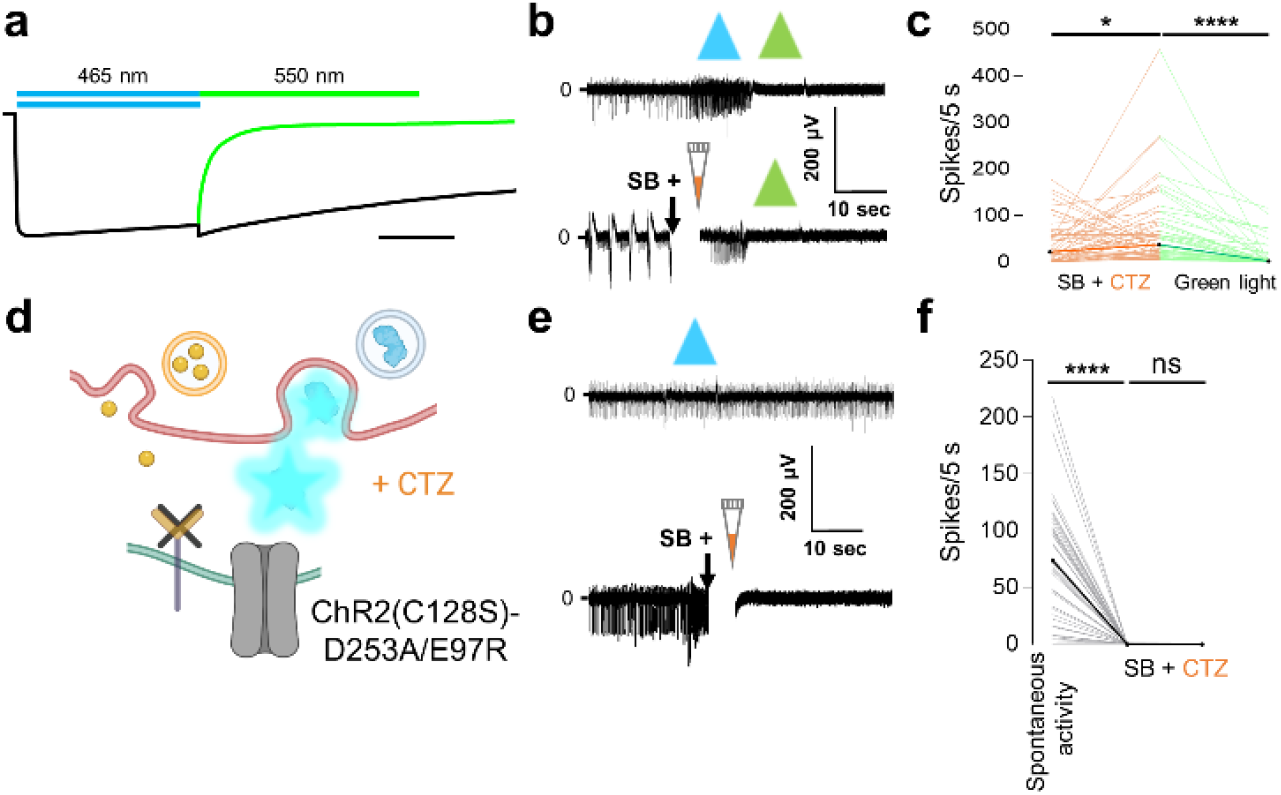
Interluminescence is mediated by bioluminescence activation of the opsin. **a** Schematic showing the typical photocurrent of the step function opsin ChR2(C128S) with a pulse of blue (465 nm) light (black trace under blue bar). There is prolonged depolarization after the light stimulation ends (continuation of black trace). Exposure to green (550 nm) light terminates the depolarization and returns the channel to its closed state (green trace under green bar; adapted from reference 15, Fig. 2a; scale bar indicates 10 seconds). **b** Representative traces of MEA recordings of postsynaptic ChR2(C128S) expressing neurons from a cortical-striatal co-culture. ChR2(C128S) can be activated by blue light and inactivated by green light (upper trace). In the presence of synaptic blockers (SB) depolarization caused by CTZ is also inactivated by green light (lower trace), indicating that the channelrhodopsin mediates the Interluminescence effect. **c** Ladder plots depict recordings from electrodes across populations as in (b, lower trace) (SB + CTZ, n=51, p=0.0369; Green Light, n=51, p<0.0001; Wilcoxon matched-pairs signed rank test). **d** Schematic of synapse with post-synaptic neuron expressing a non-functional opsin, ChR2(C128S)-D253A/E97R. **e** Representative traces of MEA recordings of postsynaptic ChR2(C128S)-D253A/E97R expressing neurons from a cortical-striatal co-culture. Postsynaptic neurons expressing the mutant opsin show no responses to either direct blue light stimulation (upper trace) or to CTZ application (lower trace), indicating that Interluminescence is a specific effect through the opsin. **f** Ladder plots of recordings under the conditions depicted in (e, lower trace) (Before vs after addition of SB, n=49, p<0.0001; Wilcoxon matched-pairs signed rank test; SB alone (after), n=21, v/s SB + CTZ (after: for non-functional opsin), n= 49, p=0.7870; Mann-Whitney test). The artifacts due to addition of reagents in MEAs are overlaid by a vertical white bar in the recording traces (the white gap right after addition of SB +CTZ). ns, not significant; *, p<0.05; ****, p<0.0001

### Interluminescence *in vivo*: Induction of cell-partner specific brain dynamics

An Interluminescent approach holds substantial distinct advantages for understanding behavior, as the complex processes underlying activities such as choice, memory and selective sensory processing inherently depend on cell-type specific interactions between multiple brain areas. These interactions relay specific signals and create the dynamic states that facilitate or suppress specific channels of information. Such inter-areal processing is highly dependent on the specific type of neurons engaged in each area.

A prominent example of this kind of cell-type specific dynamic is gamma oscillations, rhythmic patterns of activity (~30-100 Hz) that are predictive of successful sensory processing^21–23^, and are believed to amplify the relay of sensory neural signals. Neocortical gamma depends on recruitment of fast-spiking, parvalbumin-positive (FS/PV) interneurons, either through endogenous or artificially-applied glutamatergic drive, or by selective optogenetic activation^24–26^. Attentional gating of gamma in a given neocortical area is believed to be caused by excitatory intracortical^27,28^ or thalamocortical^29–31^ projections that recruit local FS/PV dynamics.

Given the potential utility of Interluminescence for *in vivo* studies, a crucial test is whether it can change network dynamics created by long-range, cell-type specific communication. To test *in vivo* efficacy, we expressed the ‘transmitting’ hPOMC1-26-sbGLuc in glutamatergic thalamic neurons (including VPm and POm) that target primary somatosensory neocortex (SI), including direct synaptic input to FS/PV^32,33^. The ‘receiver’ excitatory opsins (ChR2(C128S/D156A)) were expressed under PV-Cre mediated control in SI FS/PV^24^ (**Figure 7a** and **Supplementary Figure 7**; this subset of mice will be referred to as ‘Opsin (+)’). In SI superficial and granular layers, PV are nearly exclusively FS-type^24^. In a control group, we expressed hPOMC1-26-sbGLuc in thalamus but not the excitatory opsin in neocortex (Opsin (-)).

**Fig. 7.**
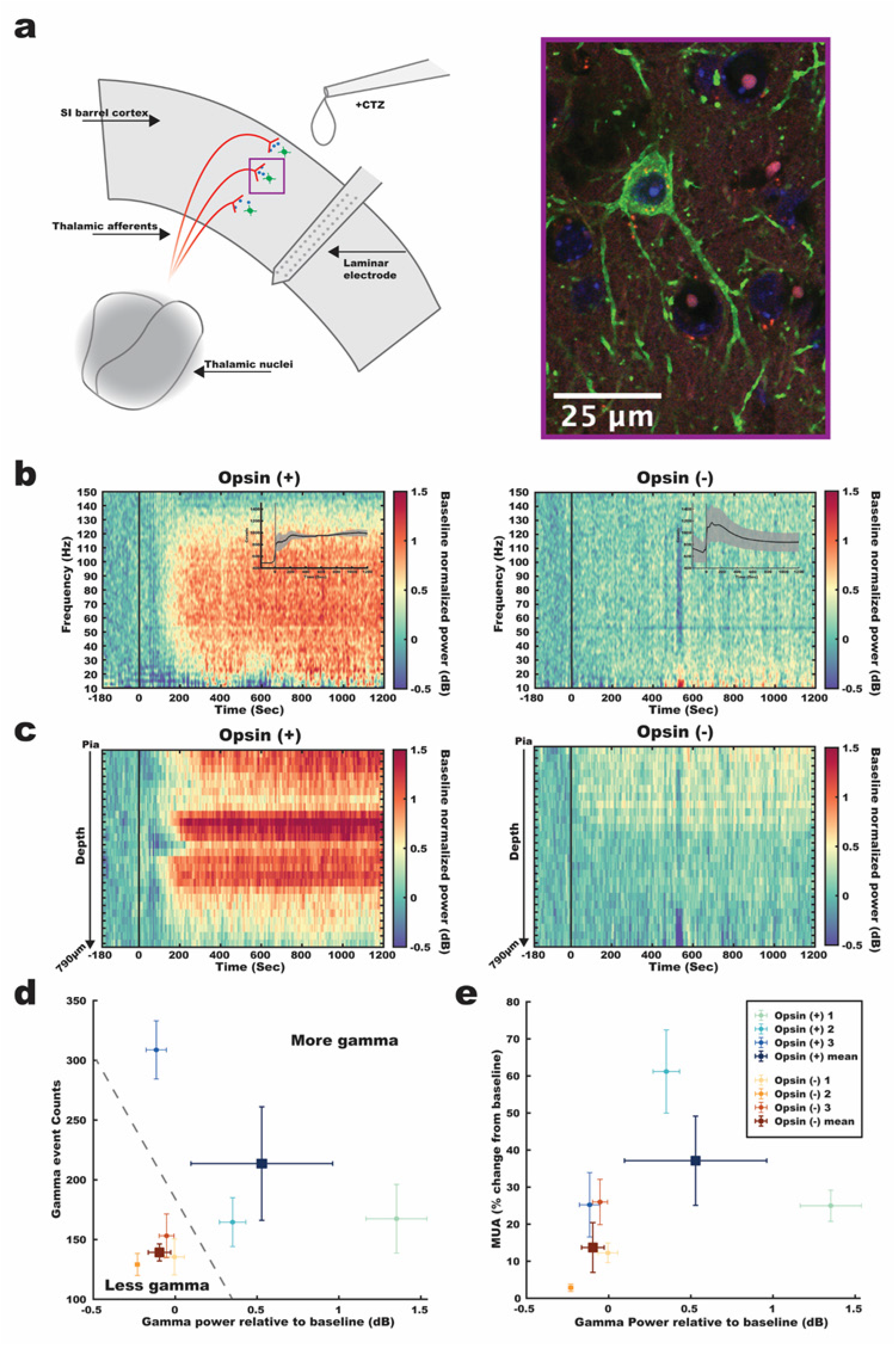
Modulation of postsynaptic neural activity by Interluminescence in vivo. **a** Schematic (left panel) of the *in vivo* Interluminescent configuration targeting somatosensory thalamic nuclei and SI Barrel cortex. Confocal image (right panel) of a PV cell expressing the excitatory step-function opsin ChR2(C128S/D156A) with an EYFP tag (green) along with thalamocortical axon terminals expressing the hPOMC1-26-sbGLuc with tdTomato tag (red) and cell nuclei stained with DAPI (blue). **b** Time-frequency spectrograms averaged across all laminar electrode contacts for the Opsin (+) (left panel) and Opsin (-) (right panel) cohorts. Time zero refers to the onset of the bioluminescent signal. The average bioluminescent signal with +/− 1 SEM in semi opaque bands is depicted as an inset for the Opsin (+) and Opsin (-) groups in the upper right corner of the respective spectrograms. The bioluminescent time series were obtained by averaging across a circular ROI adjacent to the electrode insertion point in the acquired image series. **c** Depth profiles of gamma band power (dB) (80-100Hz) across the electrode contacts for the Opsin (+) group (left panel) and Opsin (-) group (right panel). **d** Gamma band power (dB) relative to baseline plotted against gamma event counts after baseline. Small circles are individual animals, averaged across the electrode contacts. Error bars are +/− 1 SEM across the electrode contacts for a given animal. Large squares are the mean values for the Opsin (+) group and Opsin (-) group. Error bars represent +/− 1 SEM across animals. **e** Gamma band power (dB) relative to baseline plotted against MUA percent change from baseline. Same conventions as in **d**.

In Opsin (+) mice, we observed robust, broad-band gamma oscillation emergence with CTZ presentation. As shown in **Fig. 7b**, gamma initiated after CTZ presentation, consistent with the rise in bioluminescent light production (**Fig. 7b** inset). In contrast, similar changes were not observed in the Opsin (-) control group, despite robust bioluminescent photon output. These Interluminescent gamma increases were localized to more superficial and granular layers across the neocortical depth and were evident on ~half of all electrode contacts in the Opsin (+) group (**Fig. 7c:** Opsin (+) = 15.7 ± 10.1 SD electrodes/mouse; Opsin (-): M = 2.7 ± 2.5 SD).

To systematically quantify these differences, in each mouse in the Interluminescent and control groups we calculated the overall increase in gamma power on each electrode compared to baseline for the period 0-1200 seconds after bioluminescent signal onset, and the probability of significant gamma ‘events’ defined as ≥ 100ms of increased gamma (see **Methods** for details). As shown in **Fig. 7d**, the three Opsin (-) mice showed almost no increase in either overall strength or likelihood of gamma expression. In contrast, the Opsin (+) mice showed robust increases in one or both measures. Analysis of each metric at the mouse or electrode level showed significantly higher values for the Opsin (+) group (**Fig. 7d** *Mean gamma (dB) power across electrodes:* Opsin (+) Mean = 0.53 +/− 0.86 SD, Median = 0.34; Opsin (-) Mean = −0.10 +/− 0.24 SD, Median = −0.17; KS test p < .0001; 1-tailed Mann Whitney U test U = 4163, n_1_ = n_2_ = 75, p < 0.0001; *Mean gamma event number across electrodes*: Opsin (+) Mean = 213.60 +/− 139.53 SD, Median = 222; Opsin (-) Mean = 139.28 +/− 72.97 SD, Median = 149; KS test p < .0001; 1-tailed Mann Whitney U test U = 3609.50, n_1_ = n_2_ = 75, p < 0.0001; *Mean gamma power across mice* Opsin (+) Mean = 1.16 +/− 1.09 SD, Median = 0.88; Opsin (-) Mean = 0.03 +/− 0.18 SD, Median = 0.09; 1-tailed Mann Whitney U test U = 9, n_1_ = n_2_ = 3, p < 0.05; *Mean gamma event number across mice* Opsin (+) Mean = 327.25 +/− 45.58 SD, Median = 304.25; Opsin (-) Mean = 199.33 +/− 36.08 SD, Median = 203.75 ; 1-tailed Mann Whitney U test U = 9, p < .05; see **Methods** for details).

In many cases, such as during the allocation of attention, increased local action potential firing is closely tied to increases in gamma-band activity^21^. Organization of FS/PV activity into a gamma pattern is believed to enhance signal relay by creating windows of opportunity for increased local firing, and increased firing due to rebound excitation^34^. Further, increased spiking associated with gamma increases may also reflect the disproportionate contribution of FS to multi-unit activity (MUA) measures, as FS/PV firing rates are typically an order of magnitude higher than those of nearby pyramidal neurons^35,36^. That said, FS/PV evoke powerful, soma-targeted inhibition, and robust recruitment of this cell class can create local suppression of spiking^32,35^, e.g., through optogenetic FS/PV recruitment at higher light intensities^24^.

As shown in **Fig. 7e,** during CTZ presentation, Opsin (+) mice showed MUA increases from baseline that were higher than, or equivalent to, the highest mean values of Opsin (-) mice. However, only one mouse showed a significant separation from the group. Accordingly, MUA differences were significant when analyzed at the electrode, but not the mouse, level (**Fig. 7e** *Mean MUA* (*percent change from baseline*) *across electrodes*: Opsin (+) Mean = 37.14 +/− 45.50 SD, Median = 15.49; Opsin (-) Mean = 13.71 +/− 21.44 SD, Median = 8.96; KS test p < .0001; 2-tailed Mann Whitney U test U = 3899, n_1_ = n_2_ = 75, p < 0.0001; *Mean MUA* (*percent change from baseline*) *across mice*: Opsin (+) Mean = 77.72 +/− 47.40 SD, Median = 52.15; Opsin (-) Mean = 27.15 +/− 18.38 SD, Median = 27.24; 2-tailed Mann Whitney U test U = 9, p > 0.05).

## Discussion

The present study shows that Interluminescence can provide a robust, synapse-selective, activity-dependent control of neural connectivity between specific pre- and postsynaptic partners. We show that presynaptic luciferase can activate postsynaptic opsins following presynaptic activity and only if luciferin is present. This chemogenetic element of the strategy provides an additional level of experimenter control. Interluminescence is modular, as light emission from the presynaptic luciferase can, in principle, activate any postsynaptic photoreceptor, including excitatory and inhibitory opsins and light-sensing GPCRs. Further, the approach is highly specific in that Interluminescence does not seem to have a volumetric effect.

Given its features, Interluminescence has the potential to provide a platform technology, in which the activity-dependence and post-synaptic impact of Optical Synapse recruitment can be selected. In this first instantiation described here, to maximize the coupling efficiency between luciferase and opsin, we used *Gaussia* luciferase, sbGLuc, which has high light emission, and the step function opsin ChR2(C128S) and the anion channelrhodopsin 2 from *Guillardia theta*, hGtACR2 which have high photon sensitivity^6,8,9,20^. The precise photon density required to create the Interluminescent effects we observed is difficult to quantify in the abstract, as it depends on numerous factors including luciferase density, the impact of biologically specific variables in the cleft (e.g., pH sensitivity) and specific details of synaptic connectivity (e.g., synaptic distance, number of Interluminescent synapses expressing both components, location of these synapses on the post-synaptic cell, etc.). The luciferase–opsin combinations used here provide a baseline proof of concept that ample photon production was achieved in these *in vitro* and *in vivo* conditions, from a point of comparison for future luciferase-opsin pairings.

Another consideration upon moving from opsin-tethered luciferases to the two moieties positioned across the synaptic cleft was to ensure close proximity of light emitter and light sensor. In this instantiation of Interluminescence, we chose to express luciferase in synaptic vesicles, concentrating these light-producing enzymes to the presynaptic active zone, and making their release dependent on presynaptic activity. The availability of well-characterized pharmacological tools to control vesicle exocytosis, including Botulinum Toxin (BoNT), was also advantageous to confirm the dependence of Interluminescence on presynaptic vesicle fusion, a critical test of the proposed mechanism. Even though we used the POMC sorting signal to concentrate luciferase in peptide-containing LDVCs, our results suggest luciferase was present in LDCVs as well as non-peptide containing synaptic vesicles. BoNT inhibits both small synaptic vesicles (SVs) and large DCVs^37^, and immunohistochemistry revealed colocalization of luciferases in dopamine β-hydroxylase-containing and non-dopamine β-hydroxylase-containing vesicles^19^. Achieving a higher degree of specific targeting of luciferases to specific vesicles, either LDCVs or SSVs could provide a way to establish functional connectivity via Interluminescence based on the stimulation frequency. For example, there is evidence that SVs are preferentially released in response to low frequency stimuli compared to LDCVs which are preferentially released in response to higher frequency stimuli^38^. As synaptic vesicles are located in different subcellular domains of neurons, such as the cell body, dendrites, and axons, it is possible that luciferase is released at multiple sites. But, the majority of functional synaptic couplings should occur at presynaptic endings^39^ because of the requirement of close proximity of luciferase and postsynaptic opsins across a shared synaptic cleft; that is Interluminescence outside of bona fide synapses is unlikely although a possibility that could be explored in the future.

Two recently reported approaches take advantage of orthogonal neuropeptide – receptor systems for communication between genetically targeted pre- and postsynaptic partners. One is based on the insect peptide allatostatin and its receptor, both of which are inert in mammals^40^. The allatostatin receptor links via Gi/o-proteins to inhibit cyclic adenosine monophosphate (cAMP) and activate G-protein-coupled inward-rectifier potassium (GIRK) channels. Activity-dependent release of biologically active allatostatin from presynaptic neurons induces inhibition of allatostatin receptor-expressing subpopulations of postsynaptic neurons. The other system uses a *Hydra* derived presynaptically expressed neuropeptide and a matching postsynaptic cation channel that is opened by the peptide^41^. Upon activity-dependent presynaptic peptide release this heterologous synapse creates novel calcium fluxes postsynaptically, thereby activating neurons. The Interluminescence platform is complementary to these orthogonal chemical synapses and has distinct advantages. First, Interluminescence is highly modular; luciferase-emitted light can be used to activate or inhibit partnering neurons depending on the opsin expressed, an advantage over an approach that requires separate systems for activation and inhibition. Second, Interluminescence utilizes opsins as universal current conductors, effecting direct changes in membrane potential of the postsynaptic partner, an advantage over GPCR signaling pathways or Ca^2+^ flux, both of which have the potential to engage a multitude of intracellular events. Third, transmission via synthetic chemical synapses is not highly restricted to presynaptic location, consistent with neuropeptide volume transmission. In contrast, luciferase-dependent light emission decays over time and luciferases that diffuse beyond the synaptic cleft are unlikely to activate postsynaptic neurons. Fourth, synthetic chemical synapses are always ‘on,’ whenever the presynaptic cell is active, and they are not under temporal control. By contrast, Interluminescence can be temporally gated by controlling luciferin availability, a feature advantageous for assessing the behavioral impact.

In sum, Interluminescence provides a unique new technology for interrogating specific neural circuits with substantial temporal and spatial control. Interluminescence can boost or down regulate synaptic efficacy at specific synapses, it can be used to bias synapse, e.g. from inhibitory to excitatory and *vice versa*, and, in principle, it can establish new functional synaptic connections for example from silent to active. With rapid advances in the available palette of luciferases and opsins, this new strategy can expand to meet a wide array of experimental needs.

## Methods

### Materials

The luciferase substrate, coelenterazine (CTZ), was purchased from NanoLight Technology (Pinetop, AZ): Coelenterazine free base, the natural form of CTZ (NanoLight # 303), was dissolved at 50 mM in NanoFuel (NanoLight # 399); CTZ was further diluted 1:50 in culture medium for a 1 mM working solution that was further diluted 1:100 when added to MEAs for a final concentration of 10 μM. The same dilutions were carried out with just NanoFuel for ‘vehicle’. Cocktail of synaptic blockers included NBQX (abcam # ab120046), D-AP5 (abcam # ab120003), Gabazine (Sigma Aldrich # S106), CGP 55845 (Sigma Aldrich # SML0594) and Strychnine (Sigma Aldrich # S0532). Botulinum Neurotoxin (BoNT/A1) was purchased from Metabiologics (Madison, Wisconsin).

### Plasmids

The coding sequence for the N-terminal tagged luciferase construct with the leader peptide (amino acids 1-26) from the human pro-opiomelanocortin gene (hPOMC1-26)^11,12^, the *Gaussia* luciferase variant sbGLuc^10^, a P2A self-cleaving peptide, and the dTomato sequence^42^ was synthesized (Genscript) and cloned into pcDNA3.1-CAG and pAAV-hSyn to generate pcDNA3.1-CAG-hPOMC1-26-sbGLuc-P2A-dTomato and pAAV-hSyn-hPOMC1-26-sbGLuc-P2A-dTomato. Removal of P2A-dTomato and replacement by the coding sequence for EGFP generated pcDNA3.1-CAG-hPOMC1-26-sbGLuc-EGFP. Generation of pcDNA3.1-CAG-ChR2(C128S)-EYFP and its non-functional mutant pcDNA-CAG-ChR2(C128S)-E97R-D253A-EYFP are described in detail in Berglund et al. 2020^20^. The coding sequence for hGtACR2-EYFP was cloned into pcDNA3.1-CAG from pFUGW-hGtACR2-EYFP (a gift from John Spudich; Addgene plasmid # 67877; RRID:Addgene_67877).

### Virus

High titer stocks of AAV2/9-hSyn-hPOMC1-26-sbGLuc-P2A-dTomato were made in-house using previously described methods for triple plasmid transfection in HEK293FT cells^8^; larger quantities were made by ViroVek.

### In Vitro

#### Primary Neurons

Primary neurons harvested from embryonic day 18 (E18) rat embryo cortex, hippocampus or striatum of both sexes were obtained from BrainBits, LLC, and processed according to the protocol provided by the company. Briefly, tissue was incubated for 10 min at 30°C in Hibernate E (minus calcium and B27 supplement; HEB, BrainBits) containing 2 mg/ml papain (BrainBits). Papain solution was removed, replaced by HEB medium, and tissue was triturated for about 1 min (90% tissue dispersal) using a 9” sterile silanized glass Pasteur pipette (BrainBits), avoiding air bubbles. Undispersed pieces were allowed to settle for 1 min before the supernatant was transferred to a sterile 15ml tube and spun at 1500 rpm for 10 min to collect the cell pellet. The pellet was resuspended in pre-warmed and equilibrated NbActiv1 medium (BrainBits) and the cells were counted by Hemocytometer using Trypan blue stain.

#### Nucleofection

Nucleofection of E18 primary rat neurons was carried out per manufacturer’s instructions (Amaxa Rat Neuron Nucleofector Kit # VPG-1003). Briefly, 1 × 10^6^ primary neurons were collected and resuspended in 100 μl of Nucleofector Solution at room temperature. The cell suspension was combined with 1 μg plasmid DNA and transferred to the nucleofection cuvette. The Nucleofector 2b Device (LONZA # AAB-1001) was used for nucleofection with Nucleofector Program “G-013”.

#### Neuron culture on MEAs

For the mixed culture set-up on MEAs, cortical neurons nucleofected with either the luciferase construct (hPOMC1-26-sbGLuc-P2A-dTomato) or the opsin construct (ChR2(C128S)-EYFP or hGtACR2-EYFP) were mixed at a 1:1 ratio and were plated on the electrode area (1 × 10^5^ cells/10 μl/well) of 1-well MEA dishes (60MEA200/30iR-Ti; Multi Channel Systems, Germany) coated with 0.1% polyethyleneimine (Sigma # P3143) and 50 μg/ml laminin (Gibco # 23017-015) in culture medium consisting of Neurobasal Medium (Gibco # 21103-049), B-27 supplement (Gibco # 17504-044), 2 mM Glutamax (Gibco # 35050-061), and 5% Fetal Calf Serum (FCS). The following day, the medium was replaced with serum-free medium (NB-Plain medium). Half of the medium was replaced with fresh NB-Plain medium every 3–4 days thereafter. For the co-culture set-up, neurons nucleofected with either the luciferase construct (hPOMC1-26-sbGLuc-P2A-dTomato) or the opsin construct (functional opsin ChR2(C128S)- EYFP or non-functional opsin ChR2(C128S)-E97R-D253A-EYFP) were plated in separate compartments of a 2 well silicon insert (Ibidi # 80209, Germany), placed on the MEA electrodes in such a way that the total number of electrodes were approximately divided equally between the two populations. Once neurons were attached, after ~18h, the insert was removed and the populations were allowed to establish synaptic connections. Half of the medium was replaced with fresh NB-Plain medium every 3–4 days.

#### MEA recordings

MEA2100-Lite-System (Multichannel Systems, Germany) was used for all MEA recordings. Consistently spiking neurons were used for recordings between DIVs 14–25 for the mixed and co-culture set-ups; only cultures showing spontaneous electrophysiological activity were used. All-*trans* retinal (R2500; Sigma-Aldrich, St. Louis, MO) was added to the culture medium to 1 μM final concentration before electrophysiological recordings. Prior to recording, all reagents were pre-warmed to 37°C. MEAs were transferred from the CO2 incubator to the heated MEA2100 head stage maintained at 37°C, and the cultures were allowed to equilibrate for 5–10 min. The headstage was situated on a microscope stage (Zeiss Observer 1) with a fluorescent light source, allowing light stimulation of cultures at different wavelength through the objective. A micropipette was used to add reagents with the reagent drop gently touching the liquid surface, creating a time-locked artifact in the recordings. Recordings were carried out with a sample rate of 10,000 Hz. After recording, the media in the wells was replaced with fresh pre-equilibrated and pre-warmed NB-Plain media, and cultures were used for another round of recording the next day. MC Rack software was used for data acquisition. All MEA analysis was done offline with MC Rack software (Multichannel Systems; RRID: SCR_014955) and NeuroExplorer (RRID: SCR_001818). Spikes were counted when the extracellular recorded signal exceeded 9 standard deviations of the baseline noise. For assessing the effects of CTZ (10μm final concentration), only electrodes displaying the expected change in spiking activity with blue light from the fluorescent light source, i.e. opsin expressing neurons, were evaluated. Pooled data was obtained from different electrodes (a) of the same culture, (b) from different cultures, and (c) over different DIVs.

#### Electrical stimulation

Electrical stimulation on MEA co-cultures was carried out using the integrated stimulus generator in the MEA head stage (MEA2100 Stimulator). Burst stimulation pattern was selected for a 100 μA current stimulus train, with inter-pulse interval of 10ms, and the pulses within this train were repeated 5 times.

#### Un-cut vs Cut experiment

Co-cultures were allowed to mature until there was synchronous firing activity across the co-culture. Effects of blue light, current stimulation, and CTZ were recorded from these synaptically connected ‘un-cut’ co-cultures. Thereafter, in the same co-cultures, inter-population connections were severed by running a piece of thin silicon like an eraser along the midline between the two populations. These ‘cut’ co-cultures, which had lost the inter-population synchronicity, were then subjected to the same treatments (blue light, current stimulation and CTZ).

#### Synaptic blockers

The cocktail of synaptic blockers (SB) (final concentrations indicated) included NBQX (10μM), D-AP5 (50μM), Gabazine (100μM), CGP55845 (100μM) and Strychnine (1μM). Aliquots were stored at −80°C and each time thawed freshly right before the start of the MEA recording. The SB cocktail was incubated at 37°C before being added to the MEA and was added gently as 10μl drop to the neuronal media in the MEA well. For recordings involving a mixture of SB cocktail with either CTZ or vehicle, CTZ or the vehicle stocks were freshly diluted with the SB cocktail to attain the final CTZ concentration of 10μM or equivalent in case of the vehicle.

#### Botulinum Neurotoxin (BoNT)

BoNT/A1 was used as a blocker for the vesicular release of the pre-synaptic luciferase. BoNT was used at 30 ng/ml for 48 h before recording experiments.

#### Confocal Imaging

For confocal microscopy nucleofected cortical neurons were grown on Poly-D-Lysine coated coverslips (Neuvitro GG-12-PDL) in 24-well dishes until DIV 21. Neurons were fixed by completely removing the media from each well and then adding 500 μL of 4% paraformaldehyde and incubating for 15 mins at RT, followed by 3 washes for 5 min each in PBS. Neurons were permeabilized by incubating in 0.1% Triton X-100 and again were washed 3 times for 5 min each in PBS. Neurons were blocked with 1% Bovine Serum Albumin (BSA) in PBST (PBS+ 0.1% Tween 20) for 1 hr, incubated for 12 hrs at 4°C with a rabbit polyclonal anti-Dopamine β Hydroxylase (DβH) antibody (Millipore Sigma, AB 1585, diluted 1:2000 in 1% BSA in PBST), then washed 3 times, for 5 min each time, with PBST. Neurons were then incubated for 1 hr at RT with Donkey anti-Rabbit IgG H&L (Alexa Fluor 594; ab150076; diluted 1:500 in 1% BSA in PBST) and washed 3 times for 5 min each in PBST. Cells were mounted in antifade mounting media (Vectashield Hardset, H-1500-10) containing DAPI and imaged with a Nikon A1 confocal laser scanning inverted microscope using a Nikon Plan Apo VC 60x/1.40 Oil DIC N2 objective (1024×1024 μm). To image sbGLuc-eGFP the optical sections were scanned with the 561 nm laser line at 60% intensity. To detect DβH^+^ dense core vesicles with Alexa Fluor 594 the 561 nm laser line at 2% intensity was utilized. Detection was done with a 450/50 filter cube for eGFP and 595/50 for Alexa Fluor 594. The raw images were exported as TIF files and analyzed with ImageJ (Rasband, W.S., ImageJ, U. S. National Institutes of Health, Bethesda, Maryland, USA, http://imagej.nih.gov/ij/)

#### Statistics and reproducibility

All analyses were performed with Prism software (GraphPad 8.2.1; San Diego, CA), which provides evaluation of the suitability of the test for the specific data set. MEA data was collected from multiple recordings within each experiment and from multiple experiments. For randomization, the plates were switched for different treatment conditions, e.g., the plate used for CTZ treatment on one day was used for vehicle treatment the next day and vice versa. Positive control blue light treatment using the fluorescent light source was done for each recording along with other treatments as the basis for selecting electrodes for analysis of opsin expressing postsynaptic neurons. All the treatments (e.g. CTZ vs vehicle) were carried out on similar DIVs (13-21 for mixed cultures; 21-26 for co-cultures) to control for age and synaptic connectivity-related variations within neuronal cultures on MEAs. For each MEA data ladder plot, n=number of electrodes were assessed. The differences in number of spikes before and after treatment were assessed for significance. Due to non-normal distribution of data, non-parametric paired Student’s t-tests (two-tails) were used. To evaluate the within-group differences, Wilcoxon matched-pairs signed rank test was used, and to evaluate across-groups differences, Mann Whitney U tests were used with significance set at p < 0.05 (ns, not significant; *, p<0.05; **, p<0.002; ***, p<0.0002; ****, p<0.0001) using 95% confidence level. Throughout the paper, the medians are highlighted for each ladder plot in the figures. The time analyzed for the number of spikes before and after treatment was 5 sec for all ladder plots, removing the time pertaining to artifacts due to addition of reagents (indicated by the white gap area in the representative recording traces as noted in the figure legends). n and p values and the type of statistical test used are described for each ladder plot in the figure legends and results.

### In Vivo

#### Animals

Six PV-Cre mice (all male; JAX stock #008069) aged 9 to 19 weeks (M = 16.10, SD = 3.72) were used. Three mice were injected with the luciferase virus (AAV2/9-hSyn-hPOMC1-26-sbGLuc-P2A-dTomato) in somatosensory thalamus along with injections of a Cre-dependent excitatory step-function opsin in SI (Opsin (+) animals). As a control, three additional mice were also injected with the luciferase virus in somatosensory thalamus, but no opsin was introduced (Opsin (-) animals). The Opsin (-) animals thus should produce light in SI due to thalamocortical projections to SI, but no optogenetic effect ought to occur because of the absence of a postsynaptic opsin. Imaging data are unavailable for one of the Opsin (+) animals due to a software malfunction. Mice were housed in a vivarium on a reversed light-dark cycle and had free access to food and water. All procedures were conducted in accordance with the guidelines of the National Institute of Health and with approval of the Animal Care and Use Committee of Brown University.

#### Surgeries and Course of Experiment

Approximately three weeks prior to the day of the experiment, each animal was anesthetized (~1% isoflurane), fitted with a steel headpost, and injected with viral constructs via burr holes made with a dental drill. Animals received an injection of 400 nl of luciferase virus (AAV2/9-hSyn-hPOMC1-26-sbGLuc-P2A-dTomato) into the somatosensory thalamus (−1.75 A/P +/− 0.05, M/L 1.575 +/− 0.175, D/V −3.4 relative to Bregma). This injection strategy targeted somatosensory thalamus broadly, likely infecting neurons in both VPM and POm. All viral injections were performed through a glass pipette fitted in a motorized injector (Stoelting Quintessential Stereotaxic Injector, QSI). The Opsin (+) animals also received additional viral injections of 200nl of the excitatory step-function opsin (pAAV-Ef1a-DIO hChR2(C128S/D156A)-EYFP; a gift from Karl Deisseroth; Addgene viral prep # 35503-AAV1; RRID:Addgene_35503) in three locations of SI equidistantly spaced around a central SI point (−1.25 A/P, 3.25 M/L) at a depth of 350 μm. All viral constructs were delivered at a rate 50 nl/min.

After 2-3 weeks of recovery, and to allow for viral expression of the constructs, experiments were conducted under isoflurane at ~1% (0.5%-2%). A dental drill was used to make a 3mm diameter circular craniotomy centered over SI (−1.25 A/P and 3.25 M/L relative to Bregma). The exposed brain remained covered in saline throughout the experiment. The animal was moved to a light tight and electrically shielded box and continued to receive anesthesia. A 32-channel probe was inserted into cortex perpendicularly to the cortical surface at a rate of ~10μm/s using a motorized micromanipulator (Siskiyou MD7700) to a depth of 795 μm or until the highest contact on the probe disappeared from view into the cortical tissue as viewed from a stereoscope. The probe was then allowed to rest in this position for ~30 min before starting the experiment. After baseline recordings of a minimum of 3 minutes, the luciferin CTZ was introduced and recordings continued for a minimum of 20 minutes. At the conclusion of the experiment, mice were euthanized with isoflurane and perfused transcardially with 4% paraformaldehyde (PFA). The brain was removed and post fixed in 4% PFA at 4° C for approximately 48 hours after perfusion. The brain was then placed in 30% sucrose at 4° C for a minimum of 36 hours before slicing. Brains were then sectioned at 50μm on a cryostat (Leica CM30505) and mounted on glass slides. Fluorescent tags in the sectioned brains were imaged on a Zeiss LSM 800 confocal microscope to verify correct viral targeting.

#### Luciferin Delivery

Water soluble coelenterazine (Nanolight #3031) was dissolved in sterile water (1μg/ml) to yield a final concentration of 2.36 mM. The solution was loaded into a 250 μl glass syringe (Hamilton #80701) fitted with a ~1 cm length of 18-gauge plastic tubing. The Hamilton syringe and tubing were placed in a motorized injector (Stoelting Quintessential Stereotaxic Injector, QSI). The tip of the plastic tubing was lowered into the pool of saline over the craniotomy using a micromanipulator until it touched the surface of the skull. The tip of the tubing was further adjusted so that it rested at a distance of ~3mm from the opening edge of the craniotomy. The luciferin CTZ was delivered by infusing 50μl of the solution into the saline over the open craniotomy at a rate of 25 μl/min.

#### Imaging and electrophysiological recordings

Electrophysiological data was acquired using an Open Ephys acquisition board (http://www.open-ephys.org/) connected via an SPI interface cable (Intan) to a 32-channel headstage (Intan). A 32-channel laminar probe (Neuronexus, A1×32-Poly2-5mm-50s-177) was connected to the headstage. The iridium electrode contacts on the probe covered a linear length of 790 μm and were arranged into two columns of 16 contacts spaced 50 μm apart. The data were acquired using the Open Ephys GUI software at a sampling rate of 30kHz and referenced to a supra-dural silver wire inserted over the right occipital cortex. Imaging data were acquired using an Andor iXon Ultra 888 EMCCD camera attached to a Navitar Zoom 6000 lens system. The data were acquired using Andor Solis data acquisition software (Andor Solis 64 bit, v4.31). The field-of-view was centered over the craniotomy and adjusted to encompass the full diameter of the craniotomy. Images (512 x 512 pixels, ~6μm^2^/pixel) were acquired continuously at an exposure length of 1s and an electron multiplication gain of 300. The data were acquired in units corresponding to the number of electrons recorded by a given pixel. A TTL pulse synchronized the recording of the imaging and electrophysiological data.

#### Electrophysiology analysis

Offline analyses of both electrophysiological and imaging data were performed in MATLAB R2020a (The Mathworks Inc.). The electrophysiological data was down sampled to 10kHz. For each recording, electrode contacts with RMS values more than three times the interquartile range above the 3^rd^ quartile or three times the interquartile range less than the 1^st^ quartile of all 32 electrode contacts were marked as errant and removed from further analyses. Across all animals a total of seven electrode contacts were marked bad, so all reported electrophysiological data are from the remaining 25 contacts. These electrode contacts were then re-referenced to the common average reference^43^.

For the time-frequency analysis of the local field potential, the data were further down sampled to 1kHz and high-pass filtered with a cutoff at 1Hz (3^rd^ order Butterworth). Spectral analysis of the time series of each electrode contact was performed using a sliding multitapered fast Fourier transform using the Chronux software package for MATLAB (version 2.11, http://chronux.org)^44^. The time-bandwidth product for the multitaper analysis was set to 3 and 5 tapers were used. Sliding windows of 10 seconds in steps of 1 second were employed to analyze the spectro-temporal evolution of the timeseries, and each 10 second window was zero padded to a total length of 2^14^ = 16384 samples. Changes in spectral power relative to baseline for each electrode contact, time window, and frequency band were represented in decibel scale as follows:

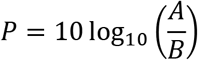

Where *B* is the average power across time in the baseline period, defined here as the 180 second period prior to the onset of bioluminescence, for a given electrode contact and frequency band, and *A* is the power in a given electrode contact, frequency bin and time bin. In addition to estimating the overall power change in the gamma band we also assessed whether high-power, but brief events in the gamma band may have increased with the introduction of the CTZ. Such events may be less detectable by the multitaper analysis due to the long time windows employed. To detect such short lived gamma events we bandpass filtered the data between 30 and 100 Hz (3^rd^ order Butterworth), Hilbert transformed the data to acquire the analytic signal and took the absolute value to acquire an estimate of the instantaneous amplitude envelope in the gamma band. Next gamma events in the post CTZ period were first defined as any data point that exceeded the 99^th^ percent jackknifed confidence interval of the baseline mean amplitude. An event was then required to exceed this threshold for at least 100 ms (i.e., at least 3 cycles at 30 Hz). Using these criteria, events were summed across the 1200 second post CTZ period for each electrode contact.

To isolate MUA, a bandpass filter (passband: 300Hz to 3000Hz, 3^rd^ order Butterworth) was applied to the data. Spikes were defined as data points less than −3 times the standard deviation, where the standard deviation was estimated as the median divided by 0.6745^18^. Spikes were then binned in increments of one second to yield a MUA timeseries for each electrode contact. Each time series was then converted to percent change from baseline.

#### Bioluminescence imaging analysis

For all images, a 3×3 pixel median filter was applied to reduce shot noise. For each animal, a circular region with a diameter of 50 pixels was placed in the region directly adjacent to the electrode shank and in front of the surface with the exposed electrode contacts. The mean of these pixels was computed for each image to yield a timeseries of bioluminescence. Since CTZ was infused into the saline over the craniotomy there was some variability in the onset of bioluminescence across animals. We were specifically interested in the relationship of bioluminescence to changes in gamma band activity and MUA, so we aligned all data to the onset of bioluminescent signal in the imaging data. We quantified the onset of bioluminescence as the peak of the discreet derivative of the bioluminescent signal. This method worked well since the bioluminescent signals in these experiments were monotonically increasing with a rapid onset.

#### Statistical analysis

As an initial descriptive statistic we calculated the number of electrode contacts that exceeded baseline for each of the three dependent measures (gamma power (dB) relative to baseline, number of gamma events and MUA percent change from baseline) in the Opsin (+) and Opsin (-) groups. A given electrode contact was considered to have exceeded baseline if its mean value in the period after bioluminescence onset was above the bootstrapped 95^th^ confidence interval of the mean of the baseline period.

For each of the three dependent measures we initially acquired three-dimensional matrices with dimensions electrode contacts, time bins, and animals. In the case of gamma power, the frequency dimension of the spectrograms was collapsed by averaging over the 30-100 Hz frequency bins. To test for significant changes in these measures we took a multi-level approach. First, for each of the dependent measures we pooled all data points across electrode contacts, time bins and animals, keeping the Opsin (+) and Opsin (-) groups separate. These pooled data sets were then submitted to a two-sample Kolmogorov-Smirnov (KS) test to test broadly for differences in the distributions of the Opsin groups. If a given KS test indicated a significant difference between the groups we then averaged across the time dimension and submitted the electrode contacts in each Opsin group to a Mann-Whitney *U* test. In the case of the gamma measures these tests were performed as one-tailed tests as we hypothesized *a priori* that upregulation of PV cells in SI via Interluminescence should increase gamma band activity. The test of MUA differences was performed as a two-tailed test as we had no *a priori* hypothesis about the directionality of these effects. The KS tests coupled with inspection of the CDFs of the two groups distributions suggested that the Opsin (+) group was positively skewed relative to the Opsin (-) group. Therefore, to sensitively test for group differences at the animal level we computed the 85^th^ percentile value for each of the measures from the array of electrode contacts for each animal. These values were then submitted to Mann-Whitney *U* tests to test for significant differences between the Opsin group at the animal level. Again, gamma band activity measures were submitted to one-tailed tests, while the MUA data were submitted to a two-tailed test.

## Acknowledgements

We thank the members of the Bioluminescence Hub (http://www.bioluminescencehub.org/) for advice and discussions. This research was supported by grants from the US National Institutes of Health (R21MH101525 to U.H.; U01NS099709 to U.H., C.I.M., N.C.S.; R01NS120832 to U.H., C. I.M., N.C.S.), the National Science Foundation (NSF NeuroNex 1707352 to C.I.M., D.L., U.H., N.C.S.), and the W.M. Keck Foundation (to C.I.M., D.L., U.H., J.A.K). M.P. was a W.M. Keck postdoctoral fellow. The figures were created with BioRender.com.

## Author Contributions

M.P. designed and performed in vitro experiments, analyzed data and wrote a draft of the paper. J.M. designed and performed in vivo experiments and analyzed data. R.S., N.F., E.L.C., A.B., A.P. participated in performing experiments. Y.B. participated in data analysis. J.A.K., N.C.S., D. L. provided critical input throughout the studies. C.I.M. and U.H. devised the Interluminescent strategy, R.S. identified the POMC vesicle targeting approach used here. U.H. and C.I.M. proposed and directed the overall study. U.H., D.L. and C.I.M. wrote the paper.

Current addresses: R.S.: The Gladstone Institutes, San Francisco, CA, USA; A.B.: Institute of Neuroscience and Physiology, University of Gothenburg, Gothenburg, Sweden; A.P.: Department of Biochemistry and Molecular Medicine, School of Medicine, UC Davis, Davis, CA, USA.; J.A.K.: Department of Psychiatry and Behavioral Sciences, Stanford University School of Medicine, Stanford, CA, USA.

## Notes

### Competing Interest Statement

The authors have declared no competing interest.

