## Supplemental Figs 1-7 for "Selective Control of Synaptically-Connected Circuit Elements by All-Optical Synapses"

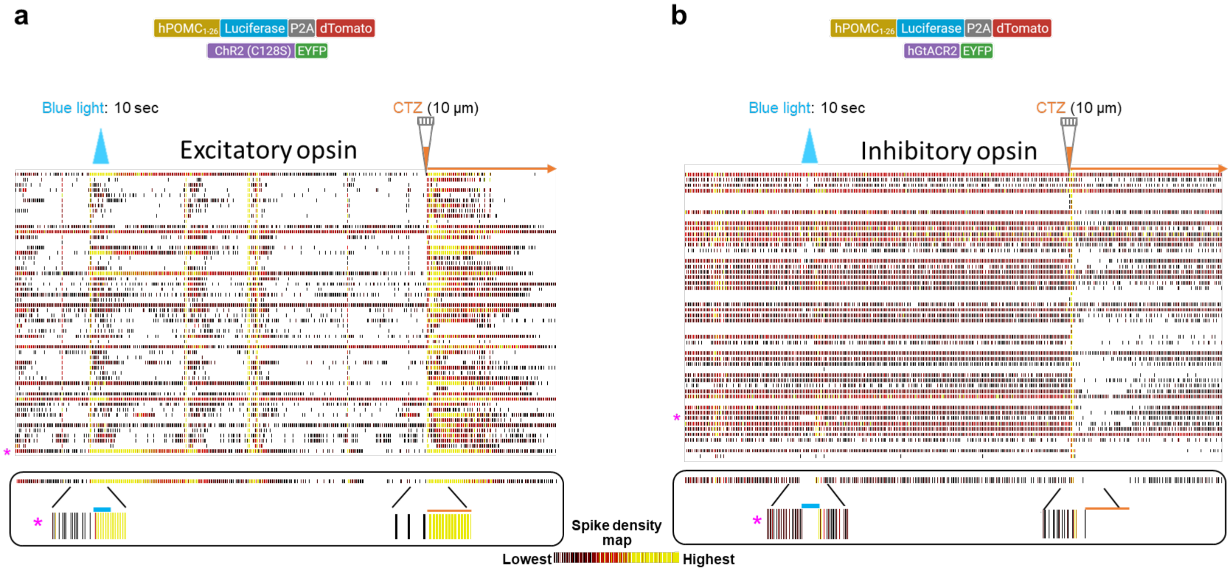


**Supplementary Figure 1. Interluminescence between pre- and postsynaptic neurons in mixed cultures.** Raster plots of representative MEA recordings showing the responses from mixed cultures of neurons expressing hPOMC1-26-sbGLuc-P2A-dTomato and neurons expressing either the excitatory opsin ChR2(C128S)-EYFP (**a**) or the inhibitory opsin hGtACR2-EYFP (**b**), respectively, to blue light and CTZ. Boxes below each plot (bottom, left and right) show a zoomed-in display of one electrode raster each marked with a magenta asterisk in the raster plot above.

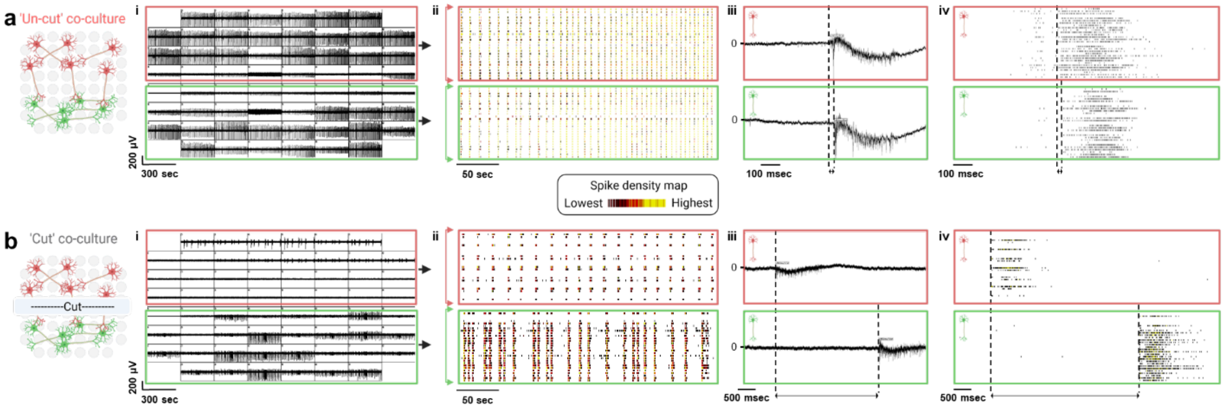


**Supplementary Figure 2. Synchrony in co-cultures of presynaptic cortical and postsynaptic hippocampal neurons on MEAs depends on intact synaptic connections. a** Illustration (left panel) showing the layout of electrodes (light grey circles) in 1-well MEAs with luciferase expressing cortical neurons over the upper half of electrodes (red) and opsin expressing hippocampal neurons over the lower half of electrodes (green) for the ‘un-cut’ co-culture. Representative example of MEA recording of a mature (DIV21) co-culture (i) showing strong synchronicity in spontaneous intra- and inter-population firing activity. Pre-synaptic and post-synaptic electrodes are outlined in red and green boxes, respectively. Raster plots (ii) from the co-culture shown in (i), demonstrating the synchronicity in firing (red arrow brackets: luciferase expressing cortical neurons from upper half of MEA; green arrow brackets: opsin expressing hippocampal neurons from lower half of MEA). Representative recording traces (iii) of individual pre- and post-synaptic electrodes, showing a very small delay in start of the firing event of the post-synaptic electrode, characteristic of synaptic transmission. Raster plot from the entire co-culture (iv) showing overall delay between the pre- and postsynaptic populations. Delay is depicted by dual-sided arrows between the dashed vertical lines for both (iii) and (iv). **b** Illustration as in **a**, left panel, with the severed connections between the two populations (‘cut’). Representative example of MEA recording of a mature (DIV21) co-culture (i) showing synchronicity in spontaneous intra-population firing activity, but a strongly disrupted inter-population synchronicity. Raster plot (ii) from the co-culture shown in (i), demonstrating the disrupted inter-population synchronicity in firing. Representative recording traces (iii) of individual cortical- and hippocampal electrodes, showing a huge time gap before the start of the firing event of the hippocampal electrode due to the loss of synaptic transmission. Raster plot from the entire co-culture (iv) showing overall delay between the cortical and hippocampal populations. Delay is depicted by dual-sided arrows between the dashed vertical lines for both (iii) and (iv).

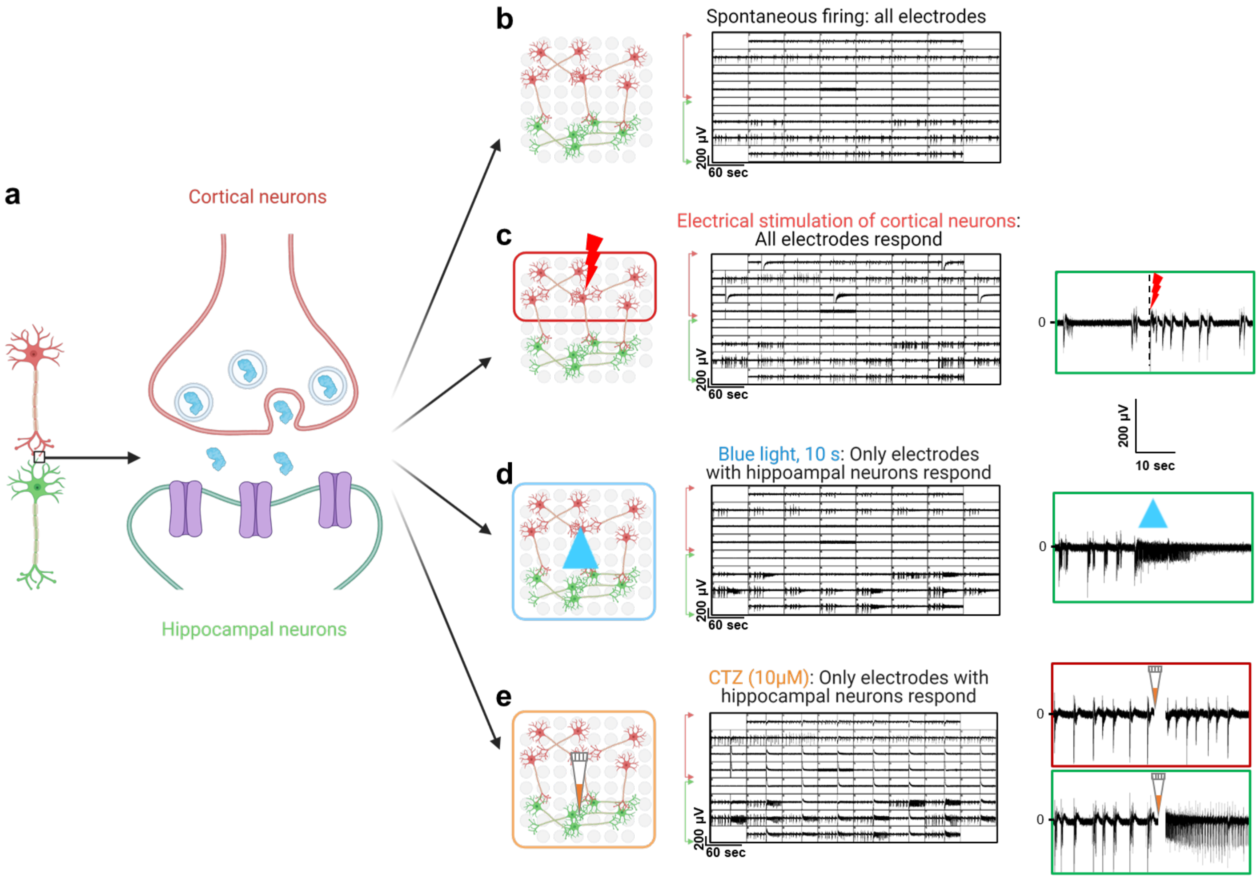


**Supplementary Figure 3. Postsynaptic activity can be elicited with presynaptic electrical stimulation and with presynaptic bioluminescence through CTZ application. a** Illustration showing luciferase expressing presynaptic cortical neuron (red) with its postsynaptic hippocampal neuron expressing the opsin (green). **b-e** Illustrations with different treatment conditions (left panels) and corresponding representative recordings from the same co-culture (right panels) for the different treatments. **b** Spontaneous and synchronous firing activity in both pre- (marked by red arrow brackets) and postsynaptic neurons (marked by green arrow brackets). **c** Electrical stimulation of cortical presynaptic neurons (red boxed electrodes: upper half of MEA) evoked a strong time-locked increase in firing activity of post-synaptic neurons (lower half of MEA). Right (green boxed): zoom-in of one example postsynaptic electrode (vertical dashed line indicates the time of presynaptic electrical stimulation shown as red bolt). **d** Exposure of the culture to blue light (blue boxed electrodes) increases activity selectively in the opsin expressing post-synaptic population. Right (green boxed): zoom-in of one example postsynaptic electrode before and with blue light stimulation. **e** Exposure of the culture to CTZ (orange boxed electrodes) increases activity selectively within the opsin expressing post-synaptic population. Right: zoom-in of one example electrode each for the luciferase expressing pre-synaptic population (red boxed) and the opsin expressing postsynaptic population (green boxed). The artifacts due to addition of reagents in MEAs are overlaid by a vertical white bar in the zoomed-in MEA recording traces (the white gap right after addition of CTZ).

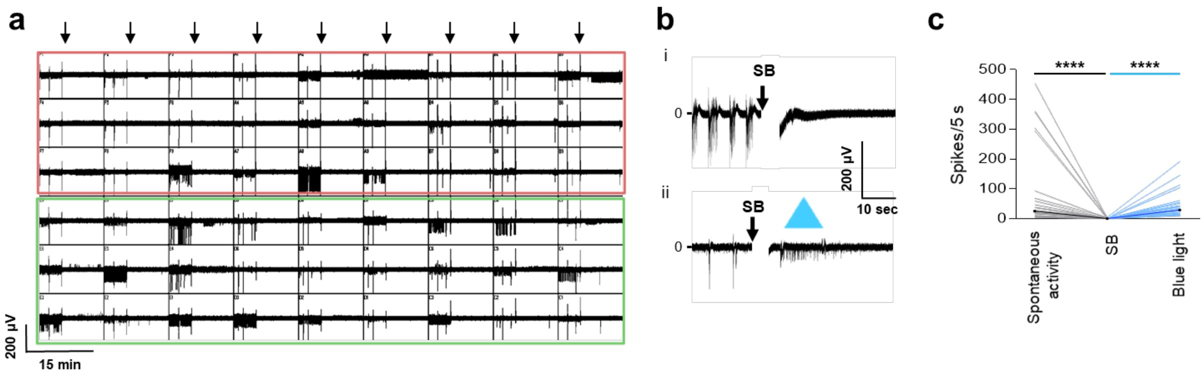


**Supplementary Figure 4. Synaptic Blockers for isolating Interluminescence effects. a** Representative example of an MEA recording from a cortical and striatal neuron co-culture with SB added about 1/3 into the recording (indicated by downwards black arrows), virtually silencing all activity in both cortical (red boxed) and striatal (green boxed) populations. **b** Zoomed-in representative traces of electrodes from striatal neurons demonstrating the silencing effect with addition of blockers (SB in i, from Fig. 4bi and ii), while preserving the ability to elicit activity in opsin expressing striatal neurons by blue light (ii). **c** Ladder plots of recordings under the conditions depicted in (b), n=27, p<0.0001; Mann-Whitney test. The artifacts due to addition of reagents in MEAs are overlaid by a vertical white bar in the recording traces (the white gap right after addition of SB). ****, p<0.0001

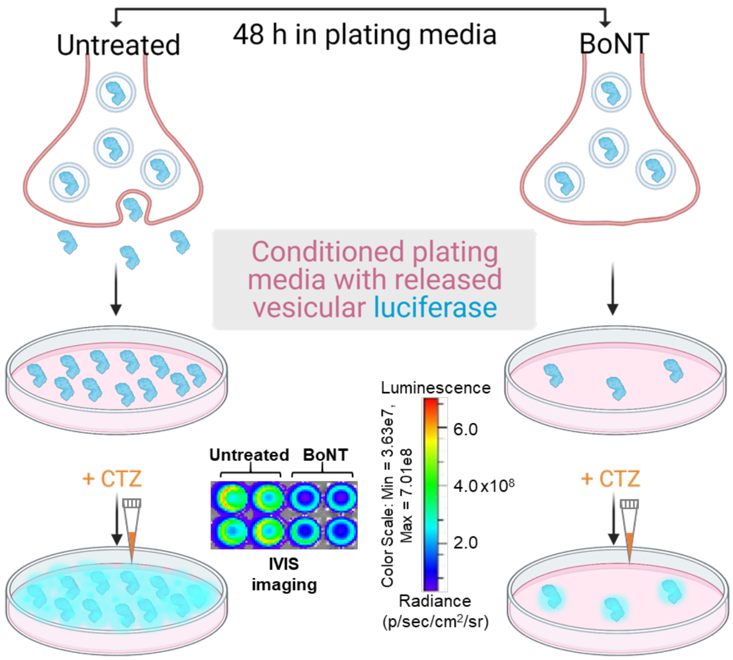


**Supplementary Figure 5. Botulinum Neurotoxin (BoNT) treatment reduces presynaptic vesicular release of luciferase**. Botulinum Neurotoxin (BoNT) treatment for 48 hours of hPOMC1-26-sbGLuc-P2A-dTomato expressing cortical neurons decreases the release of vesicular luciferase into the medium as indicated by the decrease in bioluminescence signal in ‘conditioned’ medium in IVIS images (BoNT, right side) as compared to the conditioned media from untreated luciferase expressing neurons (Untreated, left side).

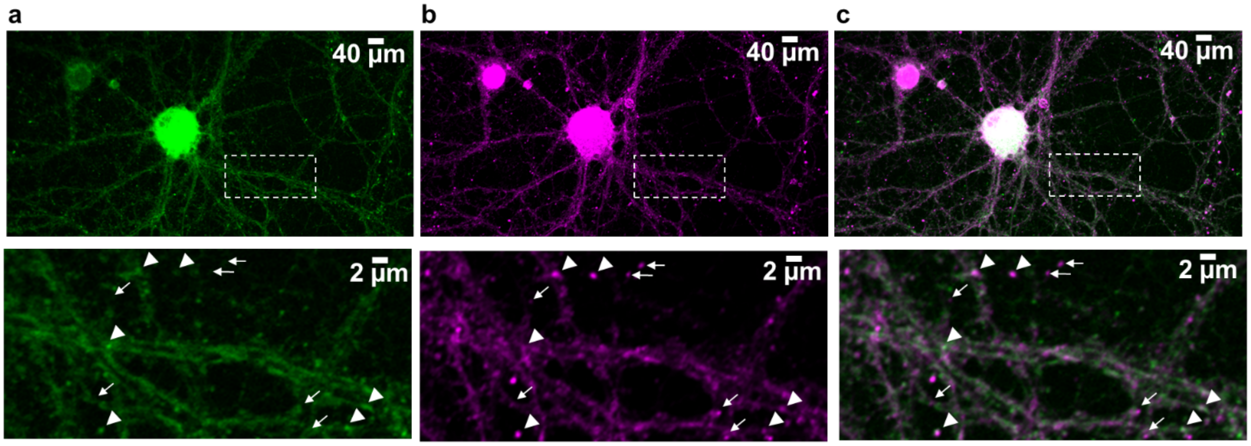


**Supplementary Figure 6. Colocalization of luciferase and dense core granule marker.** Confocal images of cortical neurons nucleofected with a plasmid encoding an hPOMC1-26-sbGLuc-eGFP fusion protein (**a**) and stained with an antibody to the dense-core granule marker dopamine β-hydroxylase (DβH) (**b**); merged images are shown in (**c**). The stippled boxes in the upper panels indicate the areas shown enlarged in the lower panels. white arrowheads, spots positive for both eGFP and DβH; white arrows, spots positive for DβH and negative for eGFP fluorescence.

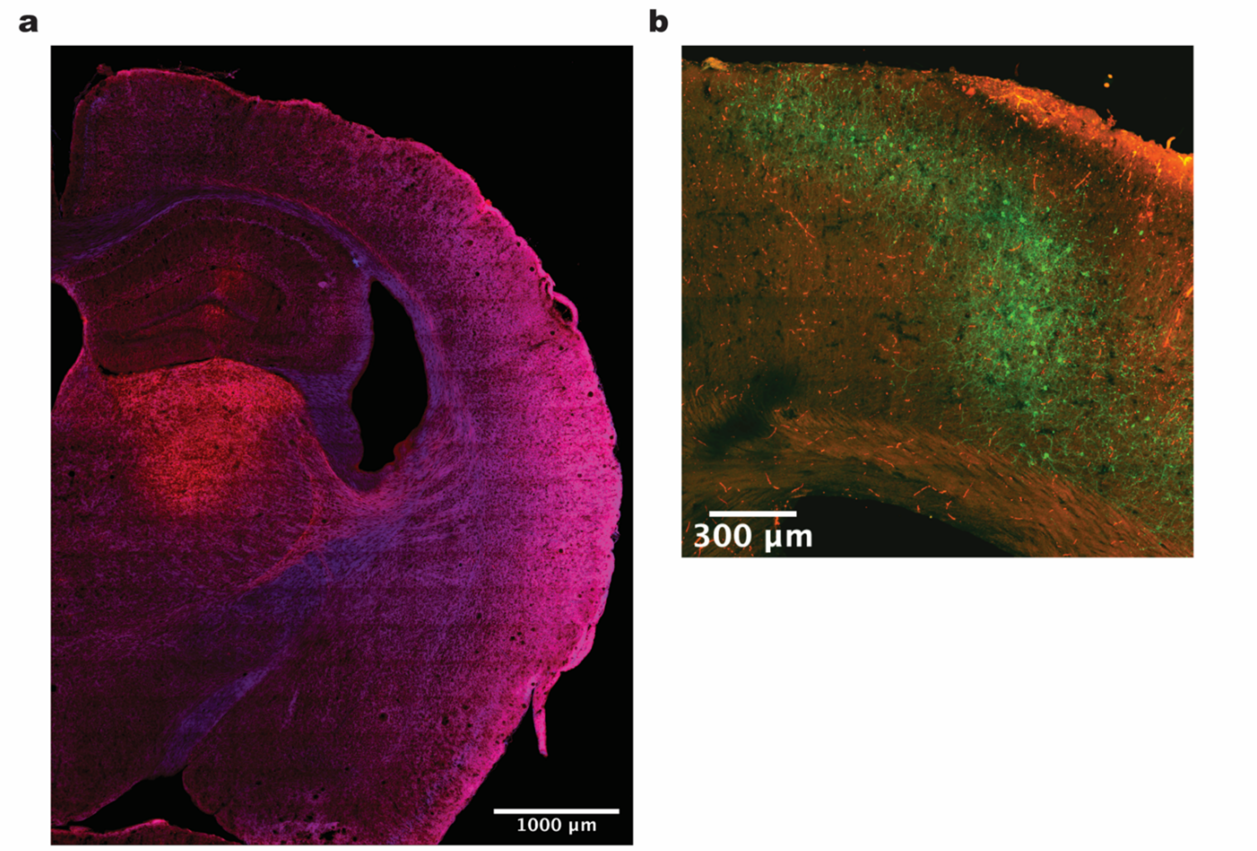


**Supplementary Figure 7. In vivo histological examples of successful viral targeting and transduction**. **a** Coronal slice from an Opsin (-) animal showing the expression of hPOMC1-26-sbGLuc with tdTomato tag (red) in thalamic nuclei merged with dapi (blue). **b** Section of barrel cortex in an Opsin (+) animal showing expression of the excitatory step-function opsin ChR2(C128S/D156A) with an EYFP tag (green) in PV cells and thalamo-cortical projecting axon collaterals and terminals expressing hPOMC1-26-sbGLuc with tdTomato tag (red).
